# Connectivity Gradient in the Human Left Inferior Frontal Gyrus: Intraoperative Cortico-Cortical Evoked Potential Study

**DOI:** 10.1101/702753

**Authors:** Takuro Nakae, Riki Matsumoto, Takeharu Kunieda, Yoshiki Arakawa, Katsuya Kobayashi, Akihiro Shimotake, Yukihiro Yamao, Takayuki Kikuchi, Toshihiko Aso, Masao Matsuhashi, Kazumichi Yoshida, Akio Ikeda, Ryosuke Takahashi, Matthew A. Lambon Ralph, Susumu Miyamoto

## Abstract

In the dual-stream model of language processing, the exact connectivity of the ventral stream to the anterior temporal lobe remains elusive. To investigate the connectivity among the inferior frontal gyrus (IFG) and the lateral part of the temporal and parietal lobes, we integrated spatiotemporal profiles of cortico-cortical evoked potentials (CCEPs) recorded intraoperatively from 14 patients who had had resective surgeries for brain tumor or epileptic focus. The 4D visualization of the combined CCEP data showed that the pars opercularis (Broca’s area) connected to the posterior temporal cortices and the supramarginal gyrus, while the pars orbitalis connected to the anterior lateral temporal cortices and the angular gyrus. Quantitative topographical analysis of CCEP connectivity confirmed an anterior-posterior gradient of connectivity from IFG stimulus sites to the temporal response sites. Reciprocality analysis indicated that the anterior part of the IFG is bi-directionally connected to the temporal or parietal area. The present study revealed that each IFG subdivision has a different connectivity to the temporal lobe with an anterior-posterior gradient and supports the classical connectivity concept of Dejerine that the frontal lobe is connected to the temporal lobe through the arcuate fasciculus and also a double-fan-shaped structure, anchored at the limen insulae.

---

Language is a unique feature of human beings that should not be impaired by surgery without the justification of clinical benefit. The posterior part of the inferior frontal gyrus (IFG), which consists of the pars opercularis (pOpe) and the pars triangularis (pTri), is known as Broca’s area, while the anterior part, the pars orbitalis (pOrb), is reported to have an executive function in semantic cognition tasks (Wagner AD et al. 2001; Gough PM et al. 2005; Hoffman P et al. 2010; Krieger-Redwood K et al. 2015). Recently, a dual-stream model of language processing has been proposed and has received wide recognition on the strength of an analogy with the processing of visual information (Hickok G and D Poeppel 2004, 2007; Saur D et al. 2008; Ueno T et al. 2011; Hickok G 2012; Gil-Robles S et al. 2013). The model consists of a dorsal stream for phonological processing and a ventral stream for semantic processing. Although this framework is generally accepted, the details of the tracts and cortices involved in the dual network remain to be established (Dick AS and P Tremblay 2012). While the superior longitudinal fasciculus (SLF) and the arcuate fasciculus (AF) are established as the main pathway of the dorsal network, additional studies are required to identify the connectivity underlying the ventral network, which presumably includes the uncinate fasciculus (UF), the inferior longitudinal fasciculus (ILF), and the inferior fronto-occipital fasciculus (IFOF).

Among these, the IFOF is reported to be involved in semantic processing based on high-frequency electrical stimulation of the white matter in awake surgery (Duffau H et al. 2005). The IFOF consists of two components, superficial and deep. The former originates from the anterior part of IFG (pOrb and pTri), while the latter originates from the dorsolateral prefrontal cortex, the middle frontal gyrus, and the orbitofrontal cortex (Sarubbo S et al. 2013). Their posterior termination includes the occipital lobe, the superior parietal lobule, and the posterior part of temporo-basal area (Martino J, C Brogna, *et al*. 2010). Duffau and coworkers reported that stimulation of the superficial component of the IFOF induced semantic errors during picture naming. The supposition that IFOF engages in semantic processing is also supported by functional studies on its cortical terminations. The anterior part of the IFG, which is part of the frontal termination of the superficial component of IFOF, has been shown to be engaged in semantic control by meta-analysis of functional magnetic resonance imaging (fMRI) and positron-emission tomographic (PET) studies (Noonan KA et al. 2013) and by interventions using transcranial magnetic stimulation (TMS) (Hoffman P et al. 2010; Jefferies E 2013). The posterior fusiform gyrus, one of the posterior terminations of IFOF, is known as the visual word-form area. A role for IFOF in semantic processing is further supported by dynamic causality modeling of BOLD (blood oxygenation level dependent) signals that showed effective connectivity from the fusiform gyrus to the anterior IFG during a semantic judgment task (Perrone-Bertolotti M et al. 2017). These pieces of evidence suggest that the superficial component of the IFOF has a semantic function. However, the anterior temporal lobe (ATL), which engages in semantic representation and is essential in semantic cognition (Spitsyna G et al. 2006; Lambon Ralph MA et al. 2010; Jefferies E 2013; Lambon Ralph MA et al. 2017), has not been reported as a posterior termination of the IFOF. ATL subregions receive the termination of the UF (temporal pole) and ILF (anterior-ventral area and temporal pole) (Binney RJ et al. 2012; Fan L et al. 2014; Egger K et al. 2015; Jung J et al. 2017; Panesar SS et al. 2018), although they seem not to be a main part of the ventral spoken language stream. Electrical stimulation of the UF does not interfere with object naming and resection of the UF is generally acceptable in neurosurgery (Duffau H et al. 2008; Duffau H et al. 2009). The ILF projects posteriorly to the occipital lobe without any frontal termination. If the superficial component of the IFOF and the anterior part of the IFG are implicated in semantic function, it would be natural to infer that the anterior temporal lobe, which is the semantic representational hub, has a direct connection to the semantic control center (the anterior part of IFG) via a subcomponent of the IFOF, although no termination of the IFOF in the anterior temporal lobe has been proven. To verify this hypothesis, we investigated in detail the connectivity between the IFG and the lateral temporal cortices.

The cortico-cortical evoked potential (CCEP) is an electrophysiological method to probe effective connectivity by applying single-pulse electrical stimulation to the cortex. The CCEPs are recorded from remote cortical areas and are supposed to reflect orthodromic propagation of the stimulus signal through the cortico-cortical connections (Matsumoto R et al. 2017; Matsumoto R and T Kunieda 2018). This method was first applied for patients with implanted subdural electrodes, which successfully delineated various functional cortical networks (Matsumoto R et al. 2004; Lacruz ME et al. 2007; Conner CR et al. 2011; Koubeissi MZ et al. 2012; Swann NC et al. 2012; Kubota Y et al. 2013; Matsuzaki N et al. 2013; Keller CJ, CJ Honey, P Megevand, et al. 2014; Enatsu R et al. 2015; Usami K et al. 2018). The connectivity pattern retrieved as CCEPs overlaps in large part with the resting-state functional connectivity measured by resting-state fMRI (Keller CJ et al. 2011; Keller CJ, CJ Honey, L Entz, *et al*. 2014). Due to its high practicality and reproducibility, CCEP has been clinically utilized to probe and monitor the connectivity of the AF during neurosurgical operations (Saito T et al. 2014; Yamao Y et al. 2014; Yamao Y et al. 2017) to ensure speech preservation (what we call “intraoperative CCEP” examination).

To investigate the patterns of connectivity from the IFG to the temporoparietal area, we systematically applied single-pulse stimulation to three IFG subdivisions (pOrb, pTri, and pOpe) and recorded CCEP responses in an intraoperative setting. Although the connectivity between the posterior IFG and the inferior parietal area and the posterior temporal area has been intensively studied (Greenlee JD et al. 2004; Matsumoto R *et al*. 2004; Greenlee JD et al. 2007; Garell PC et al. 2013; Yamao Y *et al*. 2014), this study is unique in that we visualized and analyzed the spatiotemporal dynamics of CCEP connectivity from all the IFG subdivisions, in particular from the anterior IFG, in light of the dual-stream model of language processing.

## Materials and Methods

### Participants

Fourteen patients (7 male, age 45.9 ± 17.2) were recruited for this study. The patient demographics are shown in Table 1. These patients were selected according to the following criteria from the 49 consecutive patients who underwent surgical resection of a cerebral lesion in the language-dominant hemisphere between March 2014 and July 2016. All the selected patients were supplied with the appropriate information about the study, provided written informed consent, and received intraoperative CCEP. The inclusion criteria were as follows: (1) the stimulus sites of the CCEP investigation covered all three subdivisions of the IFG (pOrb, pTri, and pOpe), which was confirmed by intraoperative photographs of the grid electrodes; (2) the recording electrodes covered the lateral temporo-parietal area; (3) no invasion or severe mass effect was observed in the temporal stem or extreme capsule, where the fibers of the ventral stream converge. See Figure 1.

**Figure 1.**
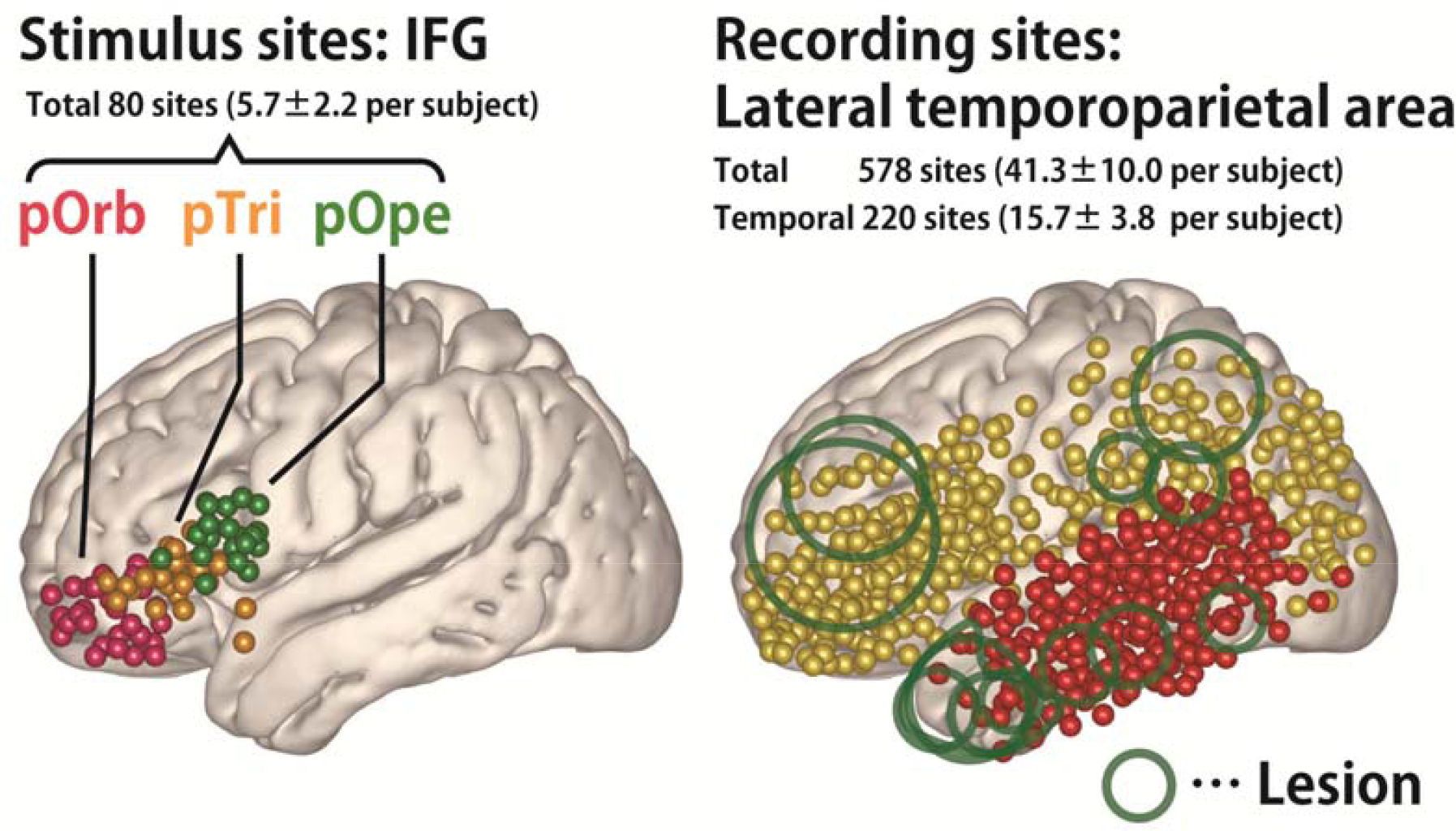
Distribution of stimulus and recording sites. **(left)** Shown are the stimulus sites in all patients, plotted in MNI (Montreal Neurological Institute) space. The coordinate of a stimulus site was defined as the midpoint of the stimulation electrode pair. **(right)** The recording sites in all patients plotted in MNI space. The red spheres indicate sites in the temporal lobe in individual brains. The green circles indicate the location of the lesions for the removal of which the patients had to undergo the surgery. The lesions are widely distributed in the frontal, parietal and temporal operculum except for the insular cortex because we excluded those patients who had a lesion or intense edema in the

**Table 1.**
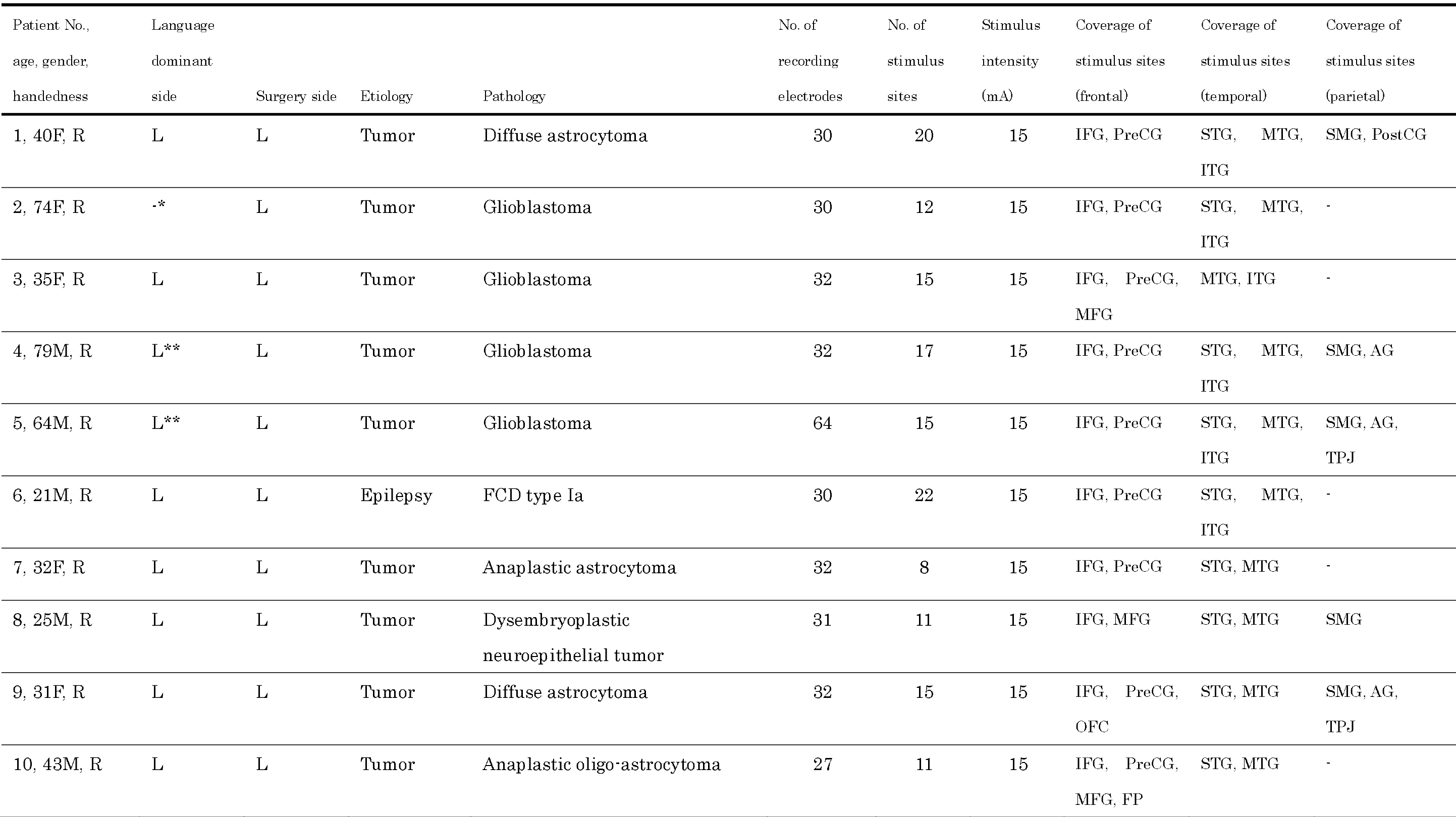
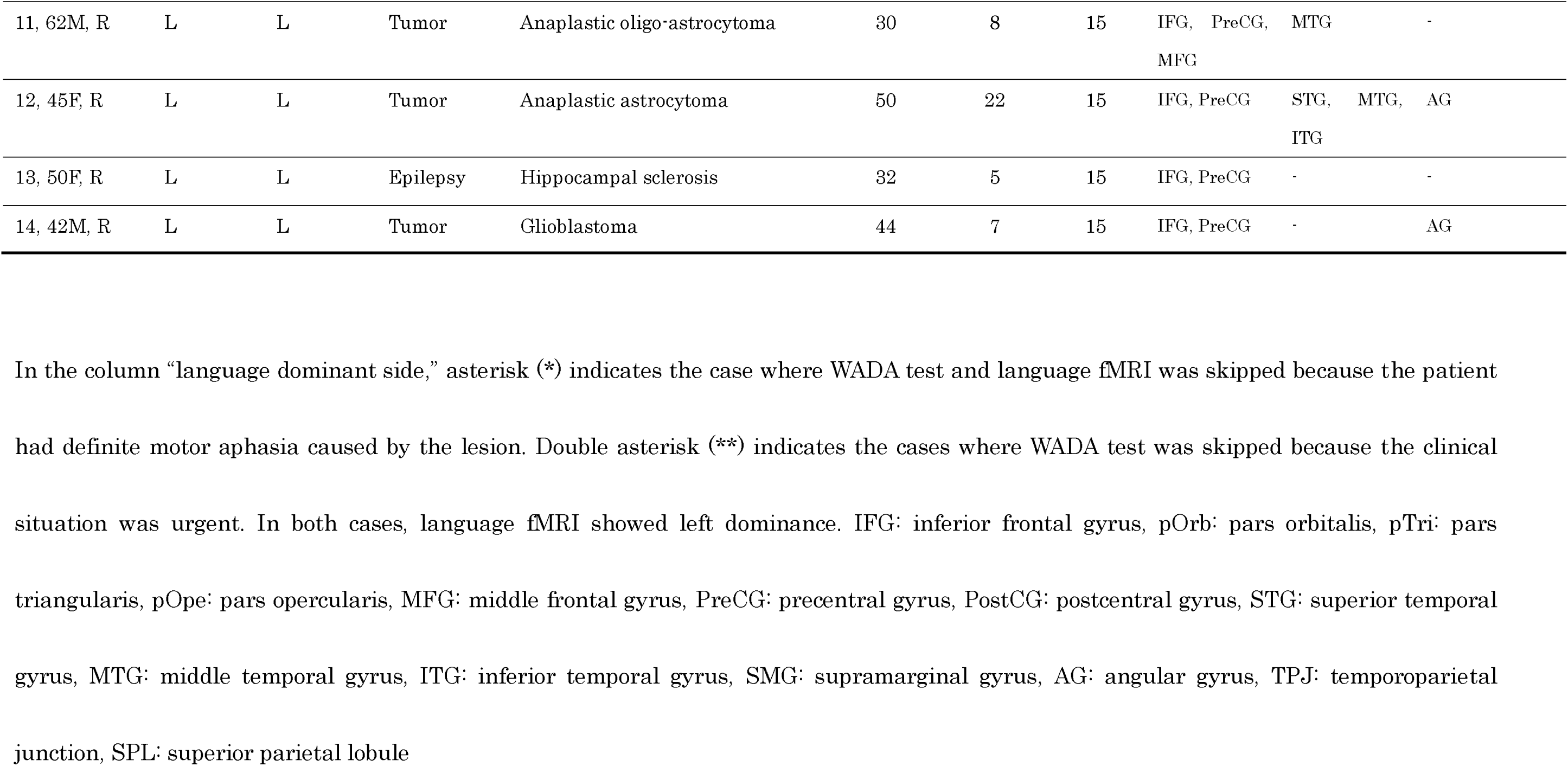
Demographic features of the patients

In 11 patients, the language-dominant hemisphere was determined by the Wada test with intra-carotid administration of propofol (Takayama M et al. 2004). In two patients (patients 4 and 5), the Wada test was omitted because of the urgent clinical situation, and language fMRI was used instead to define the dominant hemisphere (Yamao Y *et al*. 2014). One patient (patient 2) could not perform any language tasks because of motor aphasia resulting from the tumor, indicating that the affected hemisphere was language dominant. Intraoperative CCEP investigation was performed to monitor the integrity of the dorsal language pathway during surgery. The clinical usefulness of monitoring the dorsal language pathway was reported in the previous literature (Yamao Y *et al*. 2014).

This study was approved by the Ethics Committee of Kyoto University Graduate School and the Faculty of Medicine (IRB C573, C443, and C1082).

### Preparation and acquisition of intraoperative CCEP data

In all patients, the craniotomy was performed under general anesthesia. After the dura was opened, one grid electrode was placed on the frontal lobe to cover the anterior language core (Broca’s area) and another one or two grid electrodes were placed on the tempo-parietal cortices to cover the posterior language core (Wernicke’s area). The location of electrodes was always determined by clinical need (that is, monitoring of CCEP and functional mapping).

For the preoperative planning of grid placement, anatomical and functional images were acquired with a 3-Tesla magnetic-resonance scanner (Trio, Siemens, Erlangen, Germany) as described elsewhere (Yamao Y *et al*. 2014; Yamao Y *et al*. 2017). We constructed a cortical surface model from the T1-weighted image using FreeSurfer software (https://surfer.nmr.mgh.harvard.edu/).

The electrodes were made of platinum with a recording diameter of 3 mm and a center-to-center distance of 1 cm (Unique Medical Co., Ltd., Tokyo, Japan). The details of CCEP recording have been reported elsewhere (Matsumoto R *et al*. 2004; Matsumoto R et al. 2007; Matsumoto R et al. 2012; Yamao Y *et al*. 2014). A 32-channel intraoperative monitoring system (MEE 1232 Neuromaster, Nihon-Kohden, Tokyo, Japan), equipped with an electrical stimulator (MS-120B, Nihon-Kohden, Tokyo, Japan) was used for generating single pulses for stimulation, for recording the raw electrocorticogram (ECoG), and for online analysis of the averaged CCEP waveform. The raw ECoG data were recorded at a sampling rate of 5 kHz. Recordings from subdural electrodes were referenced to a scalp electrode placed on the skin over the mastoid process contralateral to the side of craniotomy.

Single-pulse electrical stimulation was applied in a bipolar fashion using a pair of adjacent electrodes. Square-wave electrical pulses of alternating polarity with a pulse width of 0.3 ms were delivered at 1 Hz. We fixed the stimulation intensity at 15 mA to shorten the investigation time since we did not have enough time to adjust the stimulus intensity in every session in the operating room. The stimulation order was as follows: First, we stimulated all possible electrode pairs in the IFG while recording the CCEP responses in the temporo-parietal area. We then stimulated selected temporo-parietal electrodes, namely, those with discrete CCEP responses to IFG stimulation, to investigate the reciprocal connectivity. All the CCEP responses analyzed in this study were recorded under general anesthesia. CCEP examination under general anesthesia was feasible since the distribution of the CCEP response did not change even though the amplitude of maximum response becomes slightly larger in the awake state (Yamao Y *et al*. 2014).

The online CCEP analysis was obtained by averaging the ECoGs (30 trials in each session) across time windows phase-locked to stimulation, each with a post-stimulus duration of 200 ms and a pre-stimulus baseline of 20 ms. We checked the reproducibility of the response in at least two sessions to distinguish the CCEP responses from baseline activities. The raw ECoG was simultaneously recorded and displayed to monitor seizure patterns during stimulation. The online CCEP analysis was used to determine the stimulation electrodes to be used for testing reciprocal connectivity. The recorded raw ECoG data were used for further offline analysis. The offline analysis was obtained by averaging ECoGs phase-locked to the stimuli (30 trials per session) with a time window of 300 ms and a baseline of 30 ms before stimulus onset.

### Visualization of the spatiotemporal dynamics of the CCEP: the 4D CCEP map

Obtaining a clear understanding of the whole connectivity pattern just from an inspection of the waveforms of individual patients is difficult because the electrode locations differ by patient. To understand the spatiotemporal dynamics based on the data derived from all patients, we created a “4D CCEP map,” that is, a 4D (time-sequence of 3D) volume image in the Montreal Neurological Institute (MNI) standard space, creating one such map for each stimulus area in the IFG (pOrb, pTri, and pOpe). For each time point (time-locked at a stimulation), all amplitude data were plotted in the MNI space, averaged and smoothed with a Gaussian kernel (see Supplementary Materials for detailed procedures.) Uneven electrode coverage was corrected by dividing the summed amplitudes by the number of measurements. To visualize the 4D representation of the CCEP, we digitally rendered a standard brain-surface model, providing each vertex with the value at the nearest-neighbor voxel in the 4D CCEP map. The time sequence of this rendered brain surface was presented as a movie. (Available in the Supplemental Data.)

### Topographical analysis of frontotemporal connectivity

To clarify the spatial relationships between the stimulus and response sites, we performed a linear regression analysis on their coordinates, which we term the “topographical analysis” of the CCEP. First, we determined the CCEP response by visual inspection of the waveforms with the following criteria:

1. The polarity is negative.
2. The amplitude is larger than 6 × the standard deviation (SD) of the baseline fluctuations. The baseline is defined as the period between 100 ms and 5 ms pre-stimulus.
3. The response is reproducible across two consecutive sessions. (30 trials are averaged for each session.)

We excluded data from electrodes located within 25 mm of the stimulus site to rule out responses due to local U-fibers since our objective was to investigate the long-range CCEP responses. Volume-conducted responses, although rare in areas > 25 mm distant, were eliminated by visual inspection because they putatively reflect large responses just under the stimulus area (Shimada S et al. 2017). We judged responses to be volume-conducted when the waveforms were almost invariant in shape and diminished steadily with distance from the stimulus site. After we inspected all the recorded waveforms to determine the early and delayed CCEP responses, the basic properties of the CCEP responses such as onset time, peak time and amplitude were stored in a database (referred to hereafter as the CCEP database) together with the MNI coordinates of the electrodes. We classified each response as early (N1) or delayed (N2) by a cluster analysis of the latency distribution (see Figure S1), although the N1 cluster determined in this method was similar to the traditional criteria of N1 (onset < 30 ms, peak < 100 ms).

Although we judged the CCEP response based solely on single waveforms, traditional waveform analysis has paid attention to locally maximal responses that seem to be the center of the response when adjacent electrodes show a similar waveform. To perform a similar analysis in this study for purposes of comparison, we identified maximum response sites in the CCEP database automatically using a MATLAB script written in-house. We defined a “max response” site as one that had the largest amplitude in the spatio-temporal neighborhood, where spatial proximity means within 15 mm of the inter-electrode distance and temporal proximity means within 5 ms of the peak time difference.

After collating the CCEP database, we investigated whether the spatial distribution of N1 responses in the temporo-parietal area differs according to the stimulus site in the IFG. Because the distribution of the response sites is parallel to the y-z plane, we verified the difference in the two-dimensional distribution (of MNI y and z coordinates) using the Wilk’s lambda test. We also evaluated the hypothesis that the more anterior the location of the stimulus site (in IFG), the more anterior is the response site (in the lateral temporal cortices). We created new coordinates for the stimulus and response sites, separately. We measured the distance between the stimulus site and the midpoint of the lower third of the precentral sulcus (see the left panel of Figure 4C) for the stimulus sites. We performed principal-component analysis for all the N1 response sites and extracted the anterior-posterior axis parallel to the temporal gyri as the first component (which we named Y1; See Figure 4C) for the response sites. The second component indicates the direction perpendicular, i.e. in a dorso-ventral temporal lobe orientation, to Y1 (named Y2). We performed linear regression analysis of X and Y1 or Y2.

### Analysis of fronto-parietal connectivity

As the number of cases that cover the parietal area is small and statistical analysis as above is not feasible, we only described the area-to-area connectivity by detecting the maximum response electrodes in each CCEP examination. We intended to perform a similar topographical analysis between the IFG and parietal lobe. Here, we focused on pOrb connectivity to the inferior parietal area as investigated by pOrb stimulation, since previous literature has reproducibly reported the CCEP connectivity between pOpe/pTri and the inferior parietal area (Matsumoto R *et al*. 2012; Entz L *et al*. 2014; Keller CJ, CJ Honey, P Megevand, *et al*. 2014; Yamao Y *et al*. 2017)

### Probing reciprocal connections from and to the pars orbitalis

We investigated the reciprocality of the connections from pOrb rather than pTri or pOpe because the latter has already been investigated in our previous reports using a different patient population (Matsumoto R *et al*. 2004; Yamao Y *et al*. 2017). We considered connectivity to be reciprocal when stimulation through the temporal (or parietal) response electrode evoked a max response in at least one of the paired stimulus electrodes in the IFG pOrb. Technical details regarding this procedure are provided as supplementary data. We calculated the reciprocality rate for six groups of stimulus-recording pairs stratified by area (fronto-temporal vs. fronto-parietal) and type of response (max response, any response, or no response). We treated the stimulus-recording pair with no CCEP response equally as those with a CCEP response in order to obtain the negative controls.

### Comparison with resting-state fMRI (rs-fMRI) connectivity

We compared the CCEP connectivity originating from the IFG pOrb with the functional connectivity revealed by the rs-fMRI for the purpose of validation. We utilized the functional connectivity maps available from the NeuroSynth website (http://neurosyngh.org/) derived from 1000 healthy subjects as a reference. For each stimulus site in IFG pOrb, a functional connectivity map was obtained from the website as a 3D volume image by specifying the stimulus coordinate as the seed voxel. We calculated the voxel-wise average of all functional connectivities obtained as described above as one volume image. We subsequently visualized the averaged connectivity map as a color map on the standard brain surface and compared it with the CCEP map by visual inspection.

## Results

### Visualization of distinct connectivity patterns from IFG subdivisions

The CCEP connectivity pattern varied distinctly when stimulation was administered through different subdivisions of the IFG. As shown in a representative case (Figure 2), the distribution of the CCEP response changed depending on whether the stimulation was applied through the pOpe, pTri or pOrb. In each patient, we observed distinct connectivity patterns for different IFG subdivisions. However, it is difficult to deduce a general rule of connectivity directly from individual cases due to limitations and variations in electrode coverage peculiar to each subject. Therefore, to systematically visualize the CCEP connectivity, we combined all patient data into a standardized map of connectivity between the IFG and the lateral temporoparietal cortices.

**Figure 2.**
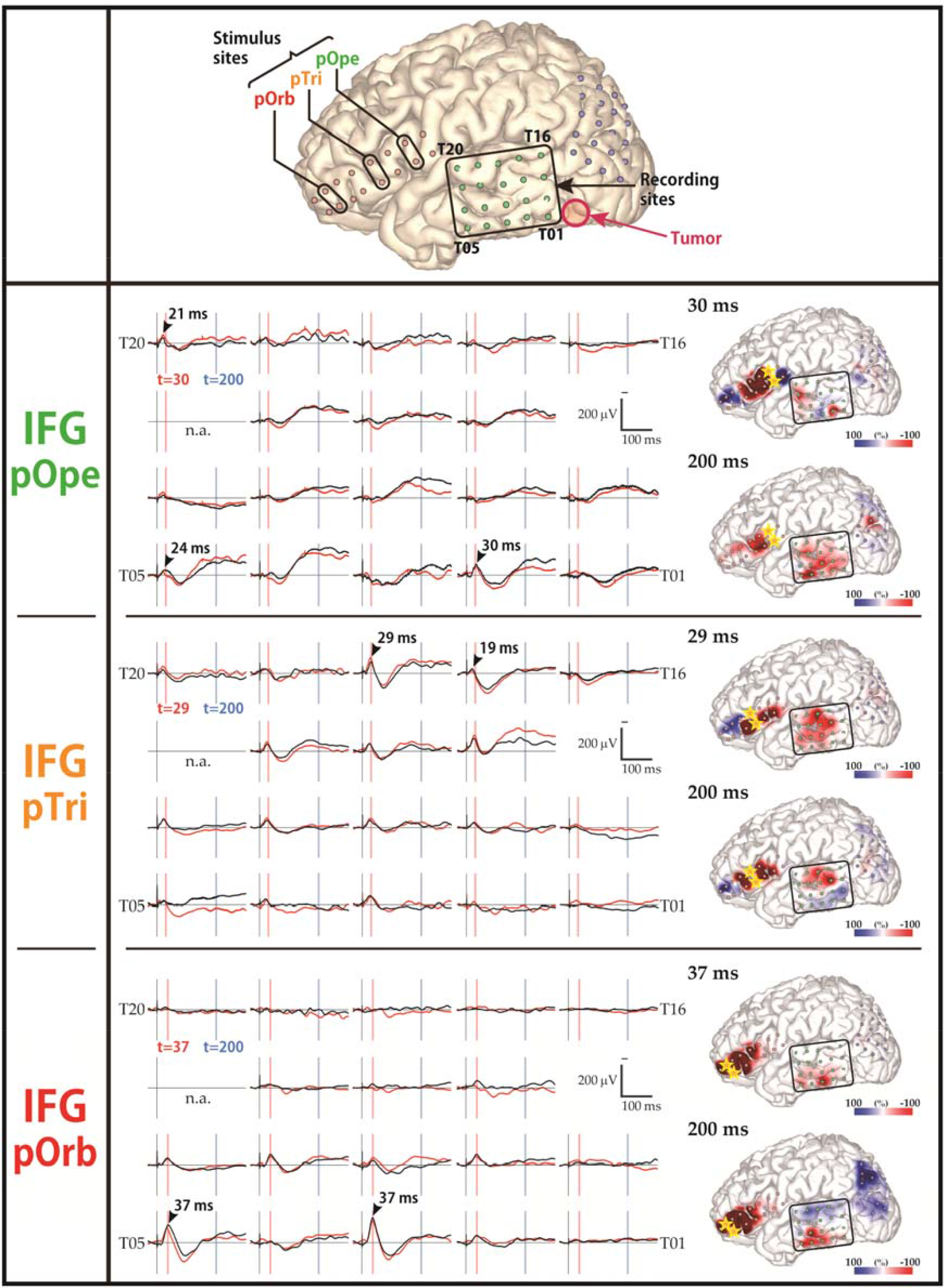
Diversity in the response pattern across stimulus sites (patient 12) **(Row 1)** Shown are the locations of all electrodes in patient 12. A 2 × 8 electrode grid was placed on the inferior frontal gyrus (IFG; contains Broca’s area) and a 4 × 5 electrode grid was placed on each of the temporal lobe and parietal lobe (jointly containing Wernicke’s area). The open red circle indicates the location of the tumor. **(Rows 2–4)** Shown are the response patterns due to stimulation in pOpe, pTri, and pOrb, respectively. pOpe, pars opercularis; pTri, pars triangularis; pOrb, pars orbitalis. Each row corresponds to one stimulus site in the IFG. **(left)** Shown are 19 of the 20 waveforms recorded by the temporal grid, with upward more negative. The lines in black and red represent the averaged waveforms of two consecutive sessions of 30 trials each. Black arrowheads indicate the “max response” sites and the peak latencies at those sites. The red and blue vertical lines indicate the timing of typical early and delayed responses, respectively. **(right)** Shown are a pair of brain-surface models painted by amplitude in the early and delayed phases, respectively. Negative amplitudes are red and positive amplitudes are blue. For each brain model, the color bar is scaled to the maximal negative response, corresponding to the right edge of the bar. Yellow stars indicate the stimulation electrodes. Electrode numbers are shown at grid corners.

Figure 3 shows the averaged response map obtained by stimulation of the three subdivisions in IFG. The 4D CCEP movie (provided as a supplemental data) demonstrates the time course of the CCEP amplitude distribution. The waveforms in Figure 3 represent the averaged temporal dynamics of the voxels included in each ROI. The center of the four spherical ROIs were located at the representative N1 (early negative) response areas in IFG stimulation: “R1” was set on the N1 response area of pOrb stimulation; “R2” and “R3,” on that of pOpe and pTri; and “R4,” on that of the stimulation in the three subdivisions. As the movie and the waveforms show, pOpe stimulation elicited prominent N1 responses in the posterior part of the temporal lobe (STG and middle temporal gyrus [MTG]) and adjacent parietal areas (supramarginal gyrus [SMG] and angular gyrus [AG]) around 30 ms after stimulus onset. After the N1 response, a larger and broader negative response (N2) with a peak latency of 150–200 ms was evoked in each response area (R2, R3, and R4). Though the averaged waveform suggests that the inferior part of the anterior lateral temporal lobe (R1) exhibited an N1 response for pOpe stimulation, we regarded this activity as a far-field potential reflecting the large response around the stimulus site; the R1 response shares the temporal dynamics of those electrodes around the stimulus site in the early phase (< 20 ms), as is demonstrated in the 4D CCEP movie. In contrast, pOrb stimulation elicited an N1 response in the anterior part of the ITG and MTG at around 40 ms after stimulation followed by a larger N2 response in the same area (see the averaged waveform of R1 in the lower panel of Figure 3). It also elicited an N1 response in the ventral part of the AG, followed by a large N2 response in the same area (see the averaged waveform of R4 in the lower panel of Figure 3). pTri stimulation showed a response pattern intermediate between those of pOpe and pOrb stimulation since its CCEP response locations comprised the posterior STG, the posterior MTG, and the AG (N1 and N2), which resemble those of pOpe stimulation, and the anterior MTG and ITG (N2), which resembles those of pOrb stimulation.

**Figure 3.**
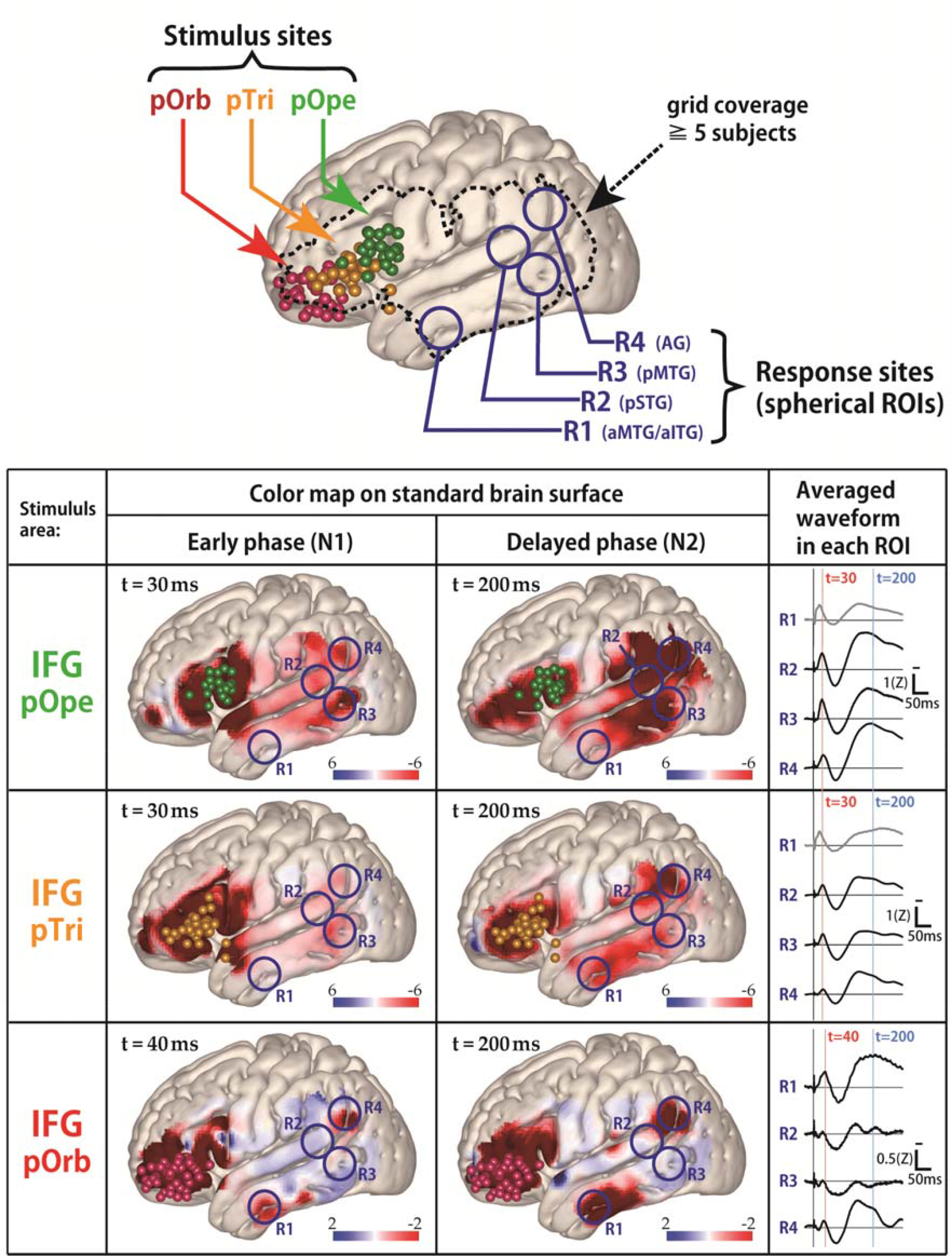
The all-patients average response map, temporal lobe. **(upper)** Shown are all stimulus sites in the IFG transferred to MNI space, stratified by subregion, and labeled with colored spheres (red, pOrb; yellow, pTri; green, pOpe). The dotted contour indicates a recording area covered by data from no fewer than 5 patients. Any voxel within the dotted contour contains the recording electrodes from no fewer than 5 patients within 15 mm of the center of the voxel. **(lower)** The table shows the averaged response maps in the early and delayed phases stratified by IFG subdivision. The 4D (movie) versions are provided in the Supplemental Materials to facilitate an intuitive understanding of the response dynamics. While stimulation of pOpe elicited a prominent response broadly in the temporoparietal area, stimulation of pOrb elicited the response in the anterior part of the medial temporal gyrus (MTG) and inferior temporal gyrus (ITG), and in the angular gyrus (AG). Stimulation of pTri elicited a pattern intermediate between that of pOpe and pOrb. We set spherical regions of interest (ROIs) of diameter of 20 mm, chosen to include the main response areas, and averaged the voxels inside each ROI to obtain the waveform for that ROI. The waveforms in the rightmost column show the time course of the response amplitudes in four ROIs located as follows: R1, in the anterior parts of the middle and inferior temporal gyri (aMTG/aITG); R2, in the posterior part of the superior temporal gyrus (pSTG); R3, in the posterior part of the middle temporal gyrus (pMTG); and R4, in the AG. Red and blue vertical lines indicate the timing of typical early and delayed responses, respectively. The R1 waveforms in the pOpe and pTri rows are grayed out because of contamination by volume conduction. (i.e., their time courses are similar to that of the stimulation neighborhood as demonstrated by the movie version of the responses.) Color bars show relative amplitude. Note that the average response map originally featured a Z-value for the amplitude, but it was smoothed by Gaussian kernel (FWHM 10 mm) for visualization. Since the Gaussian smoothing blurs the values across the adjacent voxels while keeping the sum, the values decrease after smoothing.

**Figure 4.**
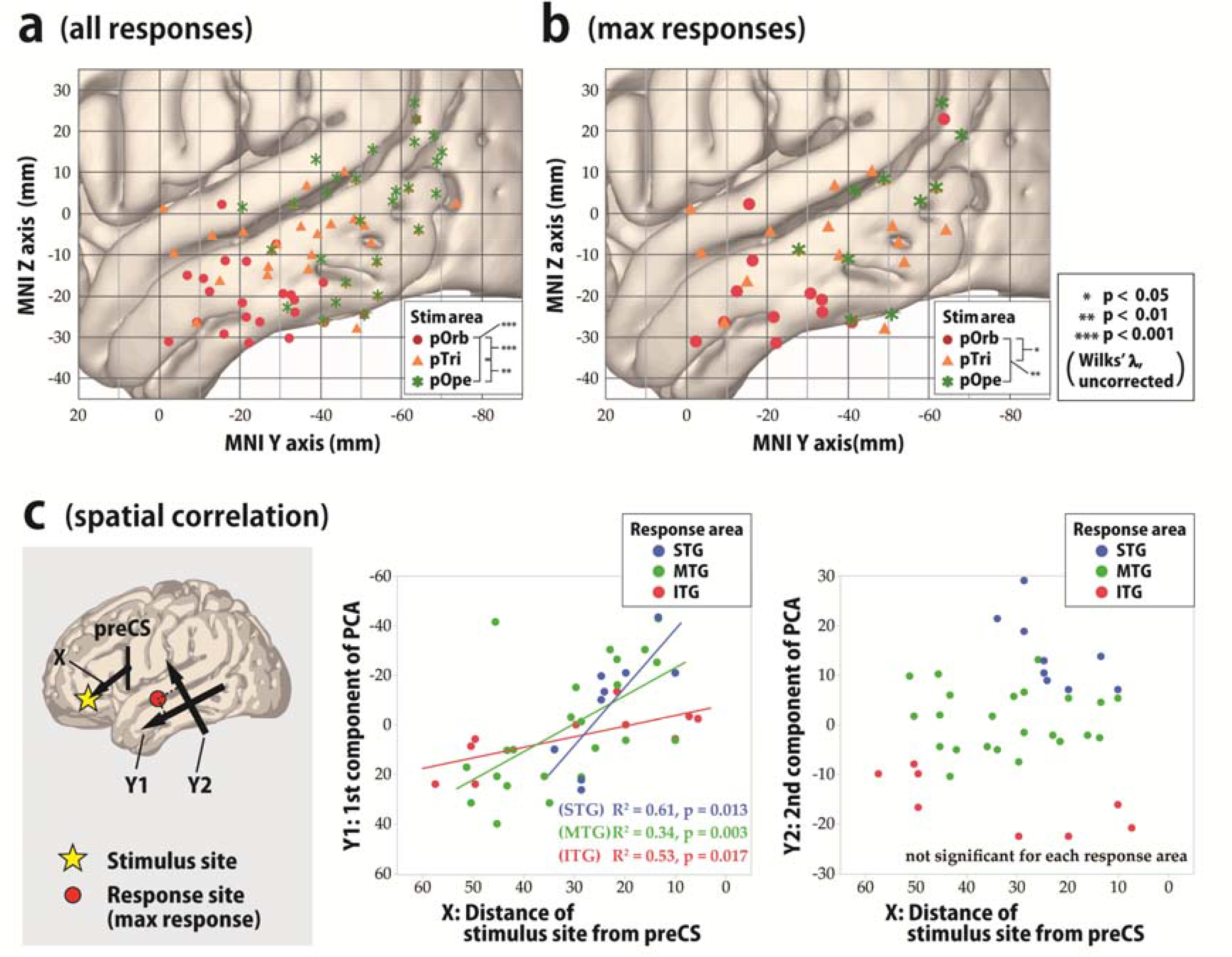
Topographical analysis of cortico-cortical evoked potential (CCEP) connectivity, temporal lobe. (A, B) To visualize the spatial distribution of the CCEP response in the lateral temporal cortices due to stimulation in the IFG, we plotted the N1 response sites in the MNI y-z plane with different colors for each stimulus area in the IFG. All response sites that satisfied the amplitude criterion of > 6 SDs of the baseline fluctuation were plotted. (A) Shown is the spatial distribution of all response sites in the lateral temporal area in the MNI y-z plane. The red circles, orange triangles, and green asterisks indicate responses to stimulation in pOrb, pTri, and pOpe, respectively. We statistically tested the difference in the response distribution for each pairwise combination of the three stimulus groups (pOrb, pTri, and pOpe) by *F*-test. We selected the stimulus sites so that the three stimulus groups would be mutually exclusive, excluding those stimulus sites whose electrodes located over the different subdivision of the IFG. (B) Shown is the spatial distribution of the maximal (max) sites. The legend for the symbols is the same as in (A). The differences among the 2D distributions were tested with Wilks’ lambda as above. (C) We generated two scatter plots to show the spatial correlation between X and Y1, and between X and Y2. In these scatter plots, linear regression analysis for each response area (STG, MTG, and ITG) was performed. Shown is the spatial correlation between the stimulus sites and the maximal response sites. The locations of the IFG stimulus sites are approximated as their MNI coordinates on the anterior-posterior axis (i.e., the distance from the midpoint of the ventral third of the precentral sulcus, PreCS). Principal component analysis (PCA) shows that the main axis of the response distribution is the anterior-posterior axis of the temporal lobe (Y1, first PCA component, **left**). Y1 is linearly correlated with the anterior-posterior position of the stimulus sites in every part of the temporal area, i.e., in the superior temporal gyrus (STG), MTG, and ITG (**middle**). Note that the value is always positive in the IFG and that a larger value indicates a more anterior location. We also used the second PCA component as the Y2-axis, which is orthogonal to the Y1-axis. Larger values on the Y2-axis indicate more superior locations. Y2 values are not correlated with the location of the stimulus site (**right**).

### Topographical analysis of frontotemporal connectivity

To statistically validate the differences in connectivity pattern visualized with the 3D average response map, we investigated the topographical distribution of the early negative (N1) responses (Figure 4A, B). In pOrb stimulation, the N1 response sites clustered in the anterior inferior part of the lateral temporal area. On the other hand, in pOpe stimulation, the N1 response sites clustered in the posterior part of the lateral temporal area. Stimulation of pTri elicited N1 responses at sites between the former two clusters. The spatial distribution was significantly different between any pair of the three parts, pOrb, pTri, and pOpe by Wilk’s lambda, *p* < 0.01. When the scatter plot was confined to the max response sites for N1, the statistical differences remained significant (Figure 4B).

The finding that pTri stimulation showed a connectivity pattern intermediate between those of pOpe and pOrb stimulation implied a gradient in the connectivity pattern revealed by IFG stimulation. We performed a linear regression analysis based on the coordinates of stimulus sites and response sites. The locations of the stimulus sites were linearly correlated with those of the N1 max response sites in the lateral temporal area (Figure 4C). In the anterior-posterior axis, the regression line could be calculated both in each temporal gyrus (STG, *Y* = 2.37 × *X* - 56.27, *R*^2^ = 0.61, *p* = 0.013; MTG, *Y* = 1.12 × *X* - 28.77, *R*^2^ = 0.34, *p* = 0.003; and ITG, *Y* = 0.43 × *X* - 3.59, *R*^2^ = 0.53, *p* = 0.017) and in the whole temporal cortex (*Y* = 0.88 × *X* - 20.29, *R*^2^ = 0.32, p < 0.0001). On the other hand, no significant correlation was observed in the superior-inferior axis. In summary, the more anterior part of the IFG connects to the more anterior part of the lateral temporal area, while the more posterior IFG connects to the more posterior temporal area, indicating a connectivity gradient along the anterior-posterior axis. We also analyzed all response sites instead of maximal response sites and observed a similar gradient of connectivity supported by linear regression analyses.

### Fronto-parietal connectivity

The averaged response map clearly demonstrated the connectivity between each IFG subdivision and the inferior parietal area. In the present study, pOrb stimulation elicited discrete parietal CCEP responses in three of five patients who had a grid on the parietal lobe (Figure 5). In all three patients, pOrb stimulation elicited early negative (N1) response in the AG, while pTri or pOpe stimulation elicited N1 response in the SMG. The N1 peak latency in the inferior parietal area was always longer in pOrb stimulation than in pTri and pOpe stimulation, as shown by the following data: patient 1, 45 ms (pOrb stimulation, AG) vs. 31 ms (pTri stimulation, SMG) and 37 ms (pTri stimulation, SMG); patient 9, 43 ms (pOrb stimulation, AG) vs. 31 ms (pTri stimulation, AG) and 34 ms (pOpe stimulation, SMG); patient 14, 34 ms (pOrb stimulation, AG) vs. 27 ms (pTri stimulation, SMG) and 31 ms (pOpe stimulation, SMG).

**Figure 5.**
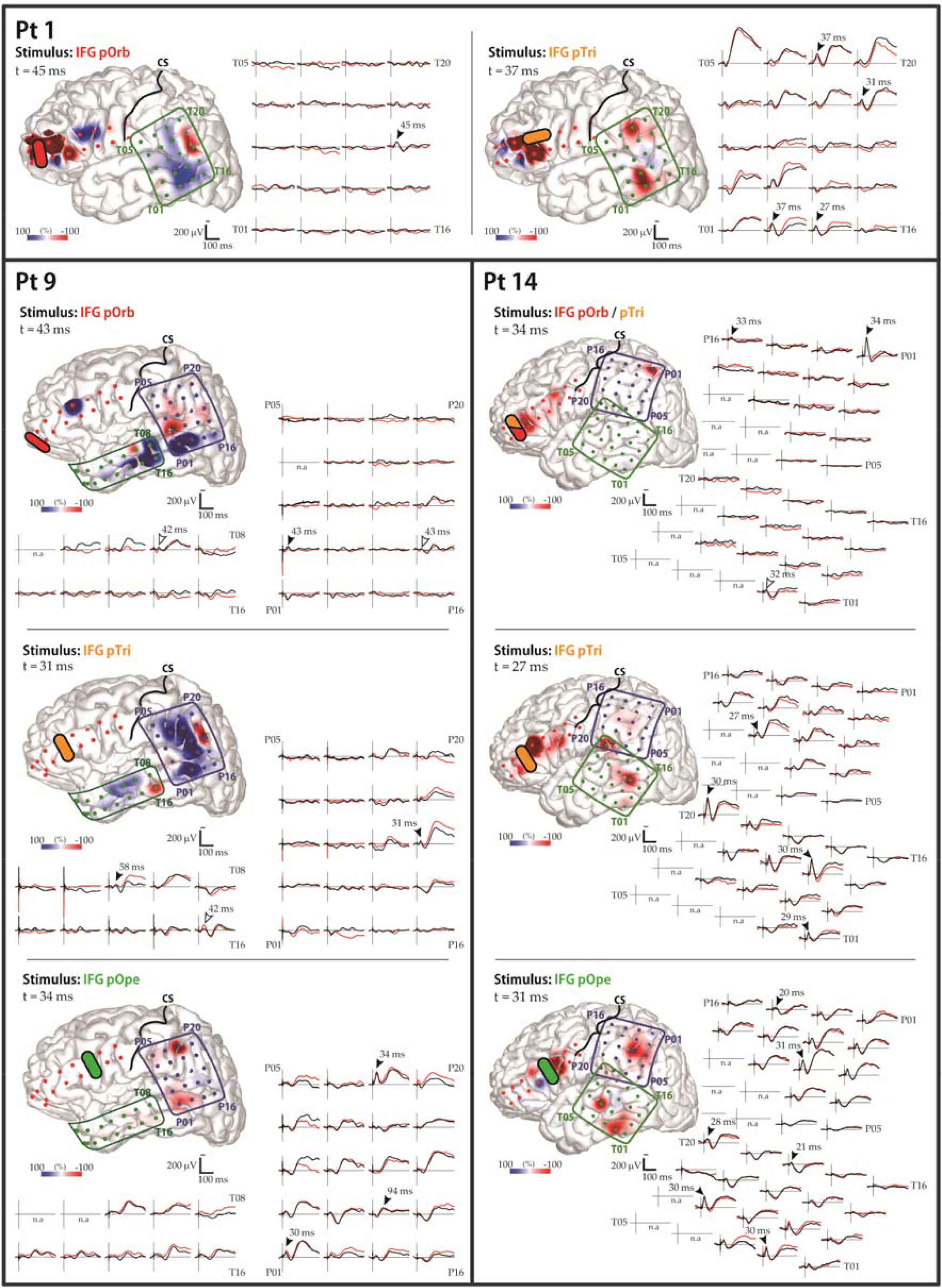
CCEP responses to IFG stimulation: Parietal lobe. Shown are the data of the three patients (1, 9, and 14) who showed frontoparietal connectivity. The waveforms in the temporal and parietal grids are shown. The black and red waveforms show the average evoked potential from two consecutive sessions. Black arrowheads indicate the maximal N1 response sites automatically extracted from the CCEP database. White arrowheads indicate maximal N1 sites added by visual inspection, which were missed by the automatic algorithm due to poor reproducibility. The brain-surface models show the spatial distribution of the early-phase response amplitudes, measured at the peak time of the parietal N1 response. The color bar is scaled to the maximal amplitudes of the parietal N1 responses in each panel. Yellow bars indicate the stimulation electrode pairs; “CS” and black lines, the central sulcus. Electrode numbers are shown at grid corners.

### Reciprocality

We investigated the occurrence rates of reciprocality in the connections between the IFG pOrb and the temporal and parietal areas (Table 2). We stimulated pOrb through a total of 41 stimulus sites and observed an anterograde CCEP response in 52 electrodes in the temporal area. Among them, 22 electrodes showed a max response. Due to limitations of time in the operating room, we were able to stimulate only 36 electrode pairs that included at least one of the anterograde response sites, and observed 25 “reciprocal” connections (25/36 = 69.44%) with max retrograde responses at the “initial” stimulus site. When the analysis is confined to max anterograde response sites, we were able to stimulate 18 electrode pairs that included at least one of the anterograde max response sites, and observed 13 reciprocal connections (13/18 = 72.22%). We performed a similar analysis of the no-response electrodes as a negative control. We aggregated all CCEP recordings that included at least one no-response electrode (333 total), and among them, we found 30 reciprocal connections (30/333 = 9.01%). The occurrence rate of reciprocal connections was significantly higher at max response sites or at all response sites than at no-response sites (unpaired *t*-test, *p* < 0.0001, uncorrected).

**Table 2.**
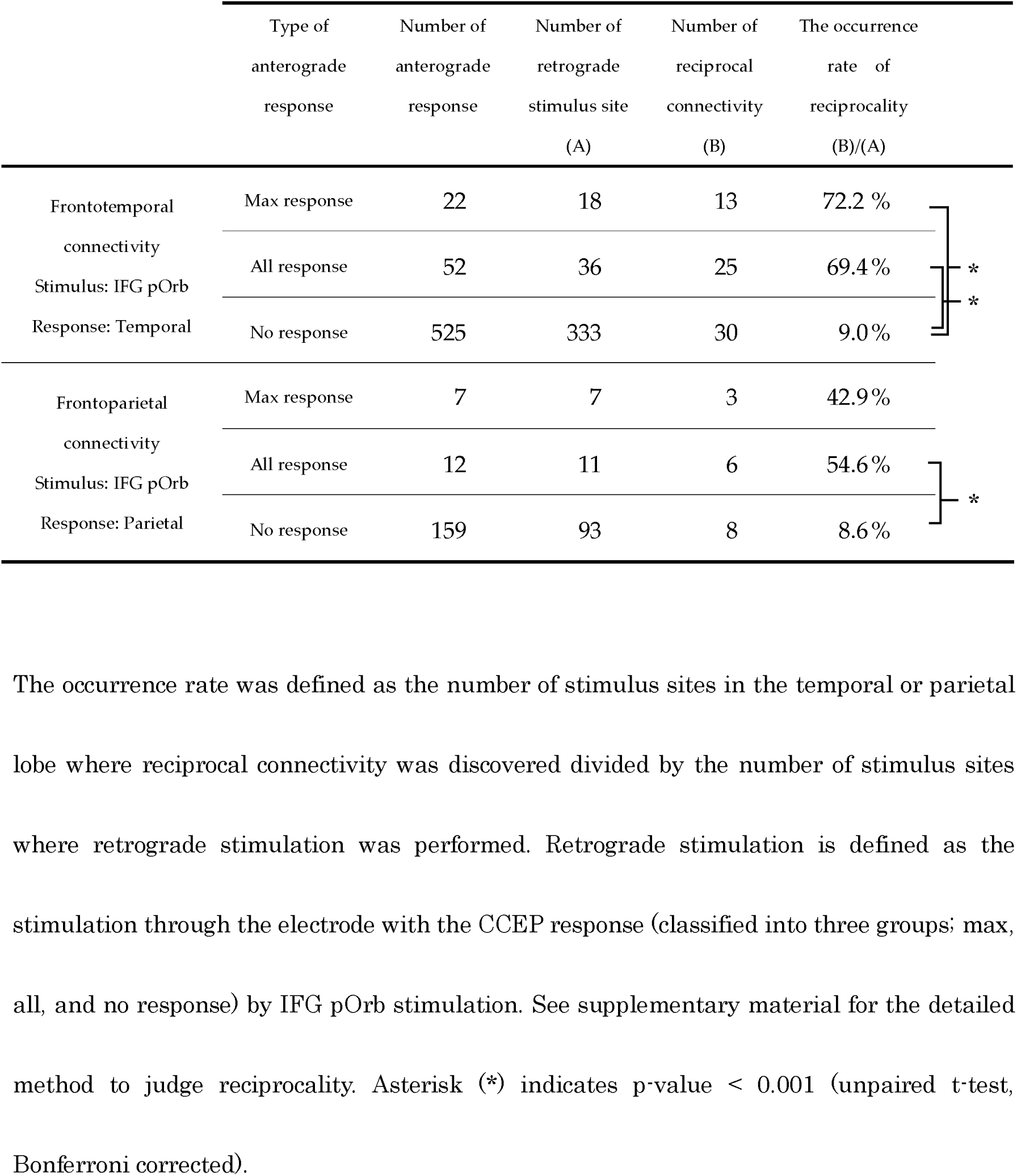
Occurrence rates of reciprocality in the connectivities originating from the IFG pOrb

We performed a similar investigation on the connectivity between pOrb and the parietal area and observed similar results, although the numbers were smaller: 3 reciprocal responses upon stimulation of 7 max anterograde response sites (3/7 = 42.86%), 6 reciprocal responses upon stimulation of 11 anterograde response sites (6/11 = 54.55%), and 8 reciprocal responses upon stimulation of 93 no-response sites (8/93 = 8.60%). The occurrence rate was significantly higher in both maximal response sites and all response sites than in no-response sites (unpaired *t*-test, *p* < 0.05, uncorrected).

### Latency and estimated conduction velocity

Table 3 shows the onset times and peak latencies of all measured waveforms. In the lateral temporal cortices, the N1 onset latency was significantly longer with pOrb stimulation than with pTri or pOpe stimulation (unpaired *t*-test, *p* < 0.005, uncorrected). The peak latency was longer with pOrb stimulation than with pOpe stimulation (unpaired *t*-test, *p* < 0.005). Similarly, in the inferior parietal lobule, pOrb stimulation showed longer latencies at onset and peak than did the other two subdivisions, although the number of available pOrb stimulations was small (*n* = 5).

**Table 3.**
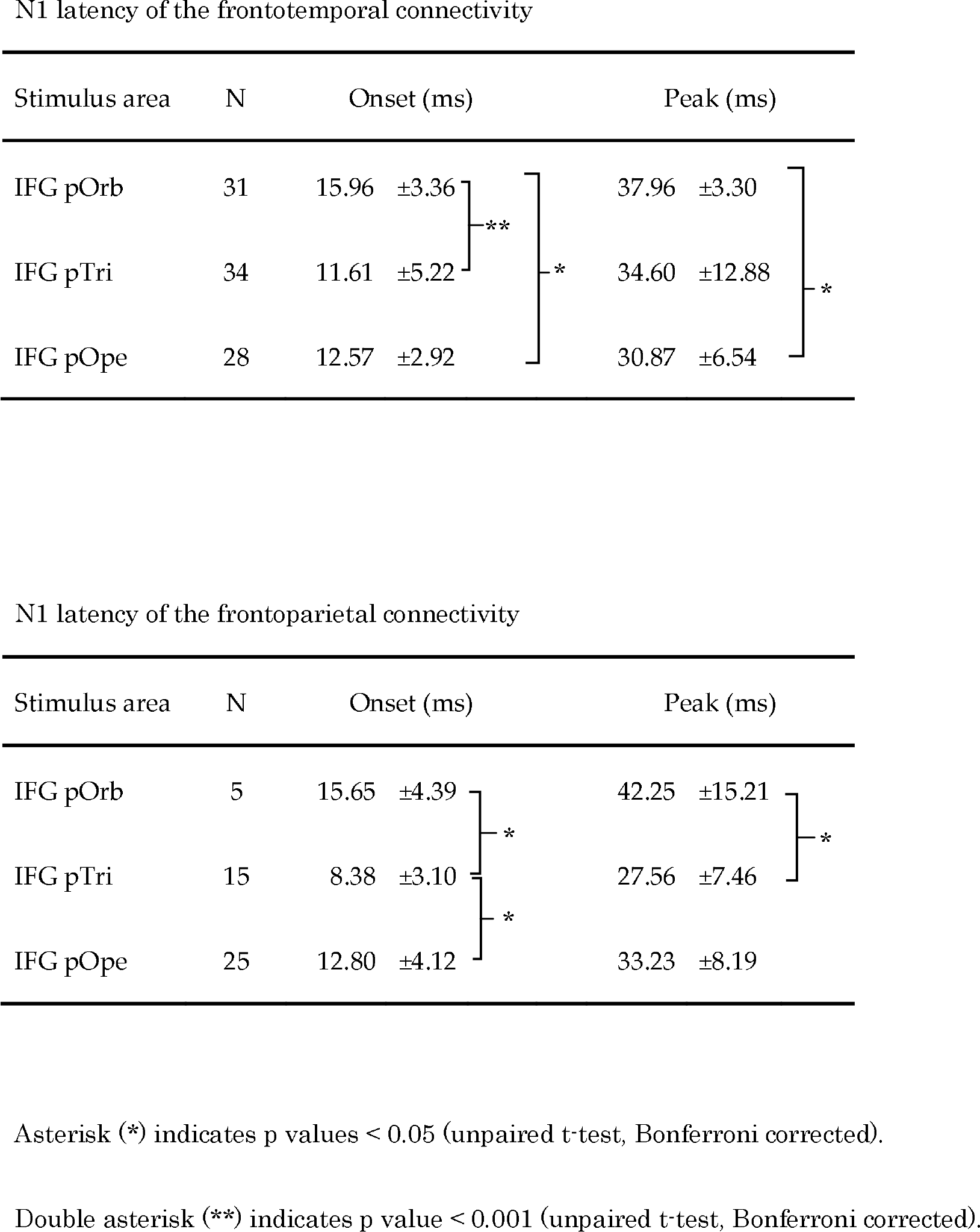
N1 Latency of the CCEP response

We plotted the onset latency vs. the Euclidean distance between the stimulus and response sites to investigate conduction velocity (see Supplemental Figure S2.) The slope of the regression line (*Y* = 0.076 × *X* + 8.2) indicated an approximate conduction velocity of 13.2 m/s (*p* < 0.05), with large variability (*R*^2^ = 0.047). We also made scatter plots for each stimulus area (pOrb, pTri, and pOpe) to compare conduction velocity among them, but no significant regression lines were found.

### Comparison with resting-state functional connectivity

The connectivity pattern elicited by stimulation of pOrb was generally similar to that of the resting-state functional connectivity obtained from the NeuroSynth database by specifying the seed as a stimulus site in IFG pOrb (Figure 6A, B). The distribution was similar between the two connectivity modalities in the anterior part of the ITG and MTG and in the inferior parietal lobule, while a difference was observed in the posterior part of the MTG (rs-fMRI positive, CCEP negative). The discrepancy is attributable to the presence of an indirect correlation via the posterior IFG (pTri and pOpe) for rs-fMRI connectivity, because pTri showed strong correlation with both pOrb and the posterior MTG. Because the resting-state functional connectivity is a measure of correlation, it inevitably visualizes a chain of strong relationships as a single indirect relationship.

**Figure 6.**
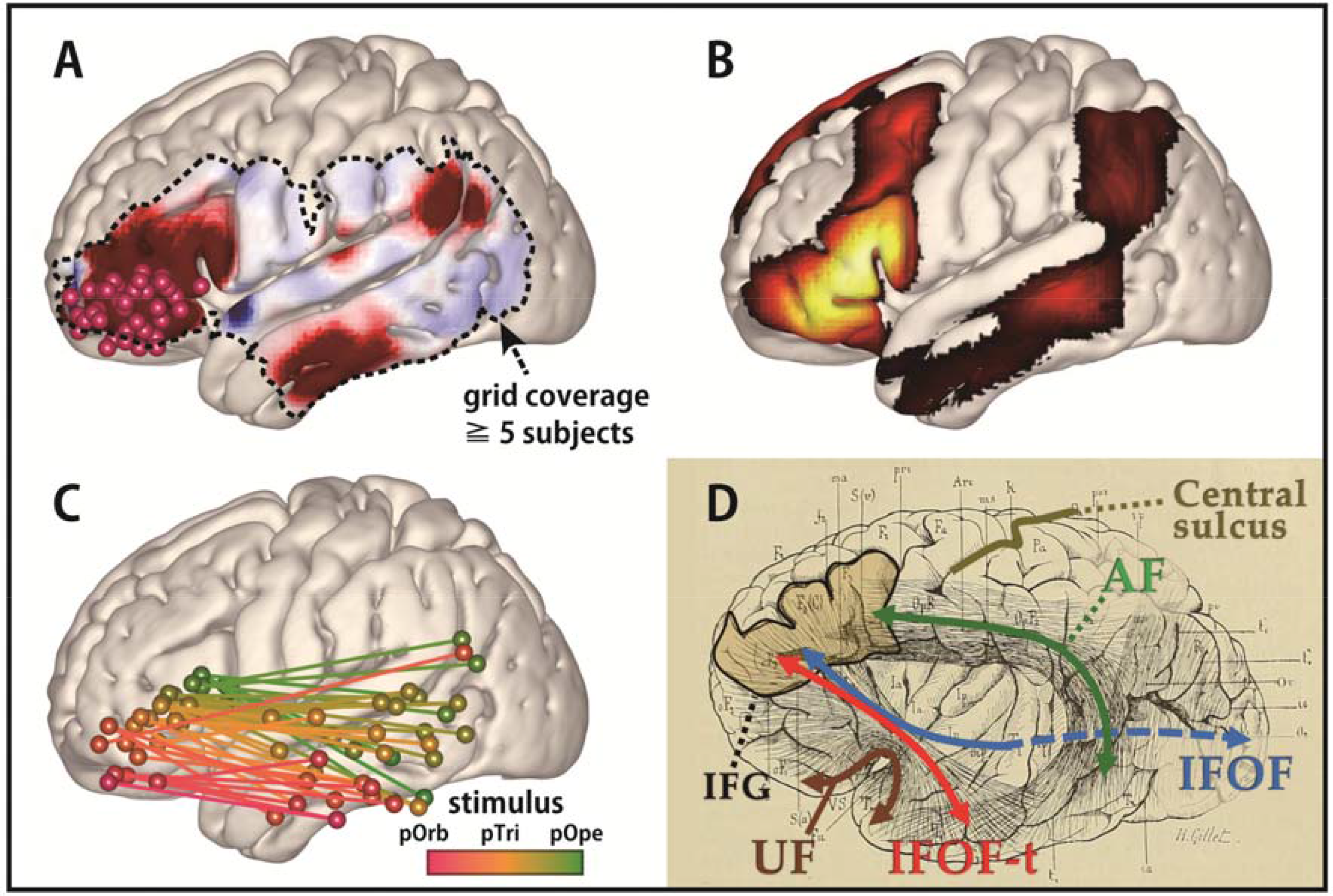
Comparison of connectivities discovered by CCEP and rs-fMRI: A connectivity gradient. (A) The averaged response map produced by stimulating pOrb, delayed phase (same as in Figure 3). (B) The functional connectivity from pOrb derived from the NeuroSynth database. In this study, the seed voxels were decided as the pOrb stimulus sites. The connectivity pattern in the lateral temporo-parietal area resembles that of CCEP. (C) The connectivity gradient between the IFG and MTG as assessed using the CCEP database. The data used in this figure are the same as that of Figure 4, although this figure shows both the stimulus and response sites in the MNI y-z plane. The color gradation indicates the anterior-posterior coordinate of the stimulus sites. The gradation from red to green corresponds to the transition from anterior to the posterior stimulation site. All pairs of stimulus and max-response sites were plotted in the MNI y-z plane to illustrate the connectivity gradient. (D) An illustration of the long tracts in the vicinity of the IFG, overlaid on the reprinted schema of white matter dissection from the classical textbook “Anatomie des centres nerveux” (Dejerine J and A Dejerine-Klumpke 1895). Major pathways are annotated with colored arrows, the IFG is outlined with a black line, and the central sulcus is outlined with a beige line. A fan-shaped structure can be seen connecting the frontal lobe and the anterior temporal lobe through the

## Discussion

Based on a compilation of CCEP data, we investigated the connectivity pattern between the IFG and the temporoparietal area. The CCEP response pattern indicated a gradual transition of connectivity from stimulus sites in the posterior IFG (pOpe) to those in the anterior IFG (pOrb). Topographical analysis of stimulus and response sites confirmed the presence of a connectivity gradient between IFG and the temporal lobe along the anterior-posterior axis. In particular, the anterior part of the IFG (pOrb) showed connectivity to the anterior lateral temporal area, which has not been well delineated by frozen-dissection, although a recent study utilizing probabilistic tractography demonstrated the connectivity between the pOrb and the lateral surface of the rostral temporal lobe (Binney RJ *et al*. 2012). We discuss the functional and clinical aspects of these results below.

### Candidate white matter pathways between the anterior IFG and temporal lobe

The present CCEP findings revealed connections between the anterior IFG (pOrb) and the anterior lateral temporal lobe. Although CCEP does not provide direct evidence about the underlying white-matter pathways, recent *in vivo* and post-mortem anatomical studies utilizing diffusion tractography and frozen dissection (Klinger’s method) potentially yield some clarification of the white-matter fibers terminating in the IFG. They consistently find that the anterior part (pOrb) and the posterior part (pTri and pOpe) of the IFG receive distinct fibers. That is, pOrb receives the termination of the IFOF (especially the superficial component) and UF, while pTri and pOpe receive terminations from the SLF and AF (Catani M et al. 2005; Barrick TR et al. 2007; Glasser MF and JK Rilling 2008; Lawes INC et al. 2008; Vassal F et al. 2016). Based on these anatomical findings, it seems plausible that upon pOrb stimulation, the electrical impulse is conveyed through the IFOF or UF, rather than the SLF or AF, to the anterior lateral temporal lobe. A different connectivity pattern from each IFG subdivision was also indicated by probabilistic tractography (Anwander A et al. 2007). That study demonstrated that the connectivity signature originating from pOpe represented the AF, while that from pOrb represented the UF and IFOF. As both structures pass through the extreme capsule in the temporal stem, we expect the existence of a pathway between the IFG pOrb and the anterior lateral temporal lobe via the extreme capsule. Since the UF mainly terminates in the temporal pole, which was not covered by the electrode grids in this study, the connectivity between the IFG pOrb and the anterior lateral temporal lobe implies the existence of the temporal branch of the IFOF, which we referred to as IFOF-t. The bundle comprised of the UF and IFOF-t can be depicted as a fan-shaped structure spreading over the temporal lobe, as illustrated in the classical textbook by Dejerine J and A Dejerine-Klumpke (Figure 6D). Here, we call this structure the “frontotemporal radiation.”

Although the IFOF-t has not been found in recent frozen-dissection or tractography studies, the existence of such connectivity is supported by our finding of reciprocality for this connectivity (Table 2) and by similarity with the resting-state functional connectivity (Figure 6). The reciprocal connectivity under the discussion implies the functional relevance of the connectivity. The resemblance between CCEP connectivity and rs-fMRI connectivity validates the existence of connectivity, as the CCEP amplitude is reported to correlate with the rs-fMRI connectivity (Keller CJ, CJ Honey, L Entz, *et al*. 2014). Furthermore, in one recent study, where whole-brain deterministic tractography was performed and virtual dissection of the UF and IFOF by a novel “stem-based” approach was carried out, a fanning structure comprising the UF and IFOF was visualized, including what we call IFOF-t (Hau J, S Sarubbo, JC Houde, et al. 2016). The discrepancy between the CCEP and the frozen-dissection/tractography results is attributable to two points. One is the presence of small fiber diameters in the ventral pathway (UF and IFOF), as revealed in an electron microscopic investigation (Liewald D et al. 2014). The other is that, like all fibers running through the extreme capsule complex and temporal stem, the IFOF-t is closely bundled with many other, major long tracts (Martino J, F Vergani, et al. 2010; Peltier J et al. 2010; Ribas EC et al. 2015; Bajada CJ, B Banks, et al. 2017). In the frozen-dissection technique, the frozen white matter is peeled along the principal fiber direction, which means that small fibers running across the major direction are destroyed (Zemmoura I et al. 2016). The tractography technique is based on the direction of local water diffusivity, which represents the principal direction of fibers within the voxel, and will, therefore, neglect the small crossing fibers (Tuch DS et al. 2003; Mukherjee P et al. 2008). CCEP relies on neurophysiological measurement, and thus makes it a more sensitive method for tracing crossing fibers.

### Candidate white matter pathways between the anterior IFG and the parietal lobe

In the present study, pOrb stimulation elicited discrete responses in the parietal area, and stimulation at the response sites revealed reciprocal connections. Based on the above discussion, pOrb stimulation is assumed to propagate through the IFOF. In the parietal termination, CCEP responses were found predominantly in the AG, which is known to be one of the posterior terminations of the IFOF (Caverzasi E et al. 2014; Hau J, S Sarubbo, G Perchey, et al. 2016). We cannot exclude the possibility that anterior IFG-AG connectivity is mediated by the SLF III because this tract is also reported to project to the AG and anteriorly as far as the dorsolateral prefrontal cortex (Mars RB et al. 2011; Seghier ML 2013; Parlatini V et al. 2017). However, to the best of our knowledge, there has been no report proving that the frontal termination of the SLF III clearly includes the IFG pOrb.

### Longer latencies of CCEP with stimulation in pOrb

The relatively long onset latency of CCEP responses seen with pOrb stimulation is consistent with the existence of the IFOF-t, since, in an electron microscopic investigation, the fiber diameter of the ventral pathway (UF and IFOF) was found to be smaller than that of the dorsal pathway (SLF) (Liewald D et al. 2014) and conduction velocity is well known to increase linearly with fiber diameter (Hursh JB 1939). The longer response latency seen with pOrb stimulation is also consistent with the fact that pOrb showed a lower myelin density than pTri or pOpe in recent myelin-density mapping studies (Glasser MF and DC Van Essen 2011; Glasser MF et al. 2016). The observation of a lower myelin density supports the possibility that the axons originating in the area are less myelinated and therefore have lower conduction velocities than those from pTri or pOpe.

### A connectivity gradient in the IFG

The linear regression analysis based on the coordinates of the stimulus and response sites indicated that the IFG is connected to the lateral temporal cortex with a gradation in the anterior-posterior axis. It is not only consistent with the presence of a fan-shaped structure, but also implies a seamless transition from the dorsal stream to the ventral stream in the IFG. Recently, a functional and connectivity gradient along the anterior-posterior axis were found not only in the IFG (Hagoort P 2005; Xiang HD et al. 2010; Udden J and J Bahlmann 2012; Thiebaut de Schotten M et al. 2016) but also in the temporal lobe (Bajada CJ, RL Jackson, et al. 2017; Jackson RL et al. 2018). Interestingly, in both, the anterior part was associated with a modality-general network and the posterior part with a modality-specific network. As Figure 6C shows, our results supported the graded functional differentiation in the IFG. Although the concept of a connectivity gradient appears in previous literature, this is the first report of an anterior-posterior gradient in the temporal projection from the IFG based on an electrical tracing method. Here, again, the gradual nature of the CCEP connectivity agrees with the fanning structure illustrated in the historical textbook, since the fan-shaped bundle of lines was drawn not only in the anterior part but also in the posterior part of the IFG (Figure 6D).

### Possibility of parcellation based on the CCEP connectivity

Our CCEP data indicated not only the existence of the two functional networks in the IFG but also the possibility of parcellation based on the CCEP connectivity. Though we can find numerous reports on the connectivity-based parcellation of human brain by means of DTI (Anwander A *et al*. 2007; Cloutman LL and MA Lambon Ralph 2012) and rs-fMRI (Arslan S et al. 2018; Jackson RL *et al*. 2018; O’Muircheartaigh J and S Jbabdi 2018), to the best of our knowledge, there are no reports on that of CCEP connectivity, though there have been reports of the whole brain connectivity matrix (ROI analysis based on template) collecting individual CCEP data (Entz L *et al*. 2014; Donos C et al. 2016). As we showed at the individual level (Figure 2) and in the group average level (Figure 3), a 1 cm difference in the stimulus site resulted in a totally different connectivity pattern and the average connectivity pattern was different for each stimulus area in the IFG. Compared to the MRI-based methods, which make use of the whole brain connectivity data, the CCEP-based parcellation seems to be difficult since the spatial resolution is no better than MRI and the CCEP data is collected only beneath the implanted electrodes. However, the CCEP-based method takes advantage of CCEP, which is based on direct electrical stimulation and is free of disadvantages such as artifacts caused by edema or tumors in MRI. Especially for such an eloquent area as the IFG on the dominant side, the CCEP connectivity-based parcellation is clinically important because it enables functional mapping without requiring the patient’s conscious cooperation as is often the case with children or patients with cognitive disturbance. Although these results must be verified with a larger population, the present study proved that the parcellation of a functionally confluent area such as the IFG solely by CCEP is feasible.

### Functional implications of the connectivity determined from pOrb

The present study demonstrated that visualizing the connectivity from the anterior part of the IFG to the anterior part of the MTG/ITG is feasible by the CCEP methodology. As discussed above, both the frontotemporal and the frontoparietal connectivity from pOrb are considered to be mediated by subcomponents of the IFOF. From a functional point of view, the IFOF is reported to have a semantic function as evidenced by intraoperative electrical stimulation at subcortical white-matter sites along the IFOF (Duffau H 2005). IFOF-t also seems to be involved in a semantic function according to the following pieces of evidence at its cortical terminations. The anterior IFG, which is the frontal termination of the IFOF-t, has been revealed by means of fMRI to engage in controlled semantic retrieval (Wagner AD *et al*. 2001; Krieger-Redwood K *et al*. 2015), and TMS in this area prolonged the response latency in a synonym judgement task (Gough PM *et al*. 2005; Hoffman P *et al*. 2010). With regards to the cortical termination in the temporal lobe, a PET activation study in healthy subjects revealed the involvement of the anterior MTG and ITG in comprehension of words presented auditorily and visually (Spitsyna G *et al*. 2006) and studies using voxel-based lesion-symptom mapping in aphasic patients found associations with semantic error in the lateral anterior temporal cortex (Walker GM et al. 2011). It appears likely that the network between these two regions, namely the IFOF-t, has a role in semantic processing.

We also determined the connectivity between pOrb and the AG, although the number of patients was small (Figure 5). As mentioned previously, tractography shows that the AG is connected with the pOrb, via the parietal branch of IFOF (Caverzasi E *et al*. 2014; Hau J, S Sarubbo, G Perchey, *et al*. 2016), which was confirmed by frozen-dissection (Curran EJ 1909; Martino J, C Brogna, *et al*. 2010). Just like pOrb, the AG is associated with a semantic role, as is proposed by meta-analysis of neuroimaging studies focusing on semantic processing (Binder JR et al. 2009), though the behavior of AG during task fMRI is significantly different from that of ATL (Humphreys GF et al. 2015), which is the semantic representational hub, as evidenced by a TMS study (Hoffman P *et al*. 2010). The facts that both the pOrb and AG are associated with semantic processing implies that the parietal branch of the IFOF may be associated with semantic processing. Even when looking outside the semantic network, a direct connection between them deserves attention because not only the AG but also the anterior IFG is involved in the default mode network (Buckner RL et al. 2008).

### Clinical implications of the CCEP examination in IFG

Previous reports from our group showed that the intraoperative CCEP with stimulation of pTri and pOpe is clinically useful to probe the posterior language area (Wernicke’s area) through AF (Yamao Y *et al*. 2014). The present study extended its clinical implication to map the whole connectivity along the anterior-posterior axis of the IFG. The graded connectivity along the IFG and the temporal lobe underlies the functional gradient in both areas as discussed above: the more anterior region connects to the more modality-general and the more posterior to the more modality-specific. This comprehensive IFG connectivity mapping would allow delineating the functional regions located in the anterior part of the IFG and temporal lobes, such as the semantic control area. In the future, we could refer to the “4D CCEP map” to guide the location of electrode placement, when more patients are enrolled to refine its quality for clinical practice.

In this study, we could not map the cortical functions in the anterior part of the IFG and the temporal lobe during awake surgeries. To investigate the semantic function, administration of specialized tasks for semantic cognition will be required. Such deliberate tasks will be more time-consuming than intraoperative tasks such as picture naming, and will demand more attention and motivation from the patients, which is difficult to achieve in intraoperative settings. Mapping studies in patients with chronically implanted electrodes for epilepsy surgery will delineate more deliberate cognitive functions in these areas. We believe mapping and preserving these higher functions out of the classical “eloquent” area such as the Broca’s area would improve the quality of life for patients undergoing neurosurgeries. In order to preserve the white matter pathway such as the temporal stem, intraoperative sequential CCEP evaluation would be clinically beneficial for patients who have lesions in the insula or the temporal stem. Detailed longitudinal neuropsychological assessments of language and semantic function warrants the functional relevance of these cortical and subcortical areas for neurosurgery.

### Study limitations and conclusion

This study investigated and clearly illustrated the connectivity between the IFG and the temporoparietal area. However, some limitations should be noted.

First, we cannot absolutely exclude the pathological effect of the lesion, though we excluded patients who had lesions or massive edema around the temporal stem, the key structure of the ventral language network.

Second, the location of the electrodes was determined by the clinical requirements of monitoring language function and the safety issues. For example, electrode grids are placed on a flat surface for stability and for keeping a distance from bridging vessels for safety. To compare the connectivity patterns among all the three subdivisions in the IFG, we included only those patients where all the three IFG subdivisions were covered by electrodes.

Third, this study lacks direct evidence on the white matter underpinning connectivity between the anterior IFG and the anterior MTG/ITG. If we observe an evoked potential in both terminals (the pOrb and the anterior MTG/ITG) by a single-pulse electrical stimulation on the white matter, it would provide proof of underpinning. Actually, we tried the stimulation of IFOF on the superior wall of the inferior horn through the removal cavity using a 1 × 4 strip electrode after anterior temporal lobectomy. However, the result was widespread CCEP responses in almost all frontal and temporal electrodes, which made interpretation difficult (unpublished data). At present, we have no direct evidence though a smaller electrode and a weaker intensity of the stimulus may improve the situation.

Fourth, this study includes no functional mapping of connectivity as mentioned in the previous section. Further studies are expected to assess the function of the connectivity observed here using electrical stimulation of the white matter and the cortices in both terminals. These assessments should be included in future studies because we believe that it is necessary for future neurosurgeons to be aware of neural structure function within the operative field even if it is out of the classical “eloquent” area.

Last, the number of patients in the present study is smaller than in other studies that visualized connectivity using CCEP. Recently, several connectivity maps based on a larger population of patients with implanted electrodes have been published (Entz L *et al*. 2014; Donos C *et al*. 2016; Trebaul L et al. 2018). Although our study includes a relatively small number of patients, it is noteworthy that all our patients had single pulse stimulation in all subdivisions of the IFG available for the observation of differences in connectivity patterns and that all data were collected within one institute, which eliminates concerns about differences in stimulus parameters and the measurement environment.

Our intraoperative CCEP data showed that the anterior IFG is connected to the anterior MTG/ITG. Combined with prior anatomical knowledge about the frontal termination of language-related fibers, these results show that the anterior IFG has a connection with the anterior MTG/ITG through the ventral stream (referred to herein as the IFOF-t) that appears as a fan-shaped structure, here named the “fronto-temporal radiation,” together with the UF and the classical IFOF. The anterior-posterior gradient in the connectivity we observed between the IFG and the temporal area suggests the presence of a gradual transition in IFG efferents between the ventral stream and the dorsal stream.

## Acknowledgments

We thank Drs. Tamaki Kobayashi, Taku Inada, Yuki Takahashi, Sei Nishida and Rika Inano for their cooperation in the intraoperative CCEP examination. The Department of Epilepsy, Movement Disorders, and Physiology, Kyoto University Graduate School of Medicine conducts Industry-Academia Collaboration Courses, supported by Eisai Co., Ltd., Nihon Kohden Corporation, Otsuka Pharmaceutical Co., and UCB Japan Co., Ltd.

## References

Anwander A, Tittgemeyer M, von Cramon DY, Friederici AD, Knosche TR. 2007. Connectivity-Based Parcellation of Broca’s Area. Cerebral cortex (New York, NY: 1991). 17:816-825.

Arslan S, Ktena SI, Makropoulos A, Robinson EC, Rueckert D, Parisot S. 2018. Human brain mapping: A systematic comparison of parcellation methods for the human cerebral cortex. NeuroImage. 170:5–30.

Bajada CJ, Banks B, Lambon Ralph MA, Cloutman LL. 2017. Reconnecting with Joseph and Augusta Dejerine: 100 years on. Brain: a journal of neurology. 140:2752–2759.

Bajada CJ, Jackson RL, Haroon HA, Azadbakht H, Parker GJM, Lambon Ralph MA, Cloutman LL. 2017. A graded tractographic parcellation of the temporal lobe. NeuroImage. 155:503–512.

Barrick TR, Lawes IN, Mackay CE, Clark CA. 2007. White matter pathway asymmetry underlies functional lateralization. Cerebral cortex (New York, NY: 1991). 17:591–598.

Binder JR, Desai RH, Graves WW, Conant LL. 2009. Where is the semantic system? A critical review and meta-analysis of 120 functional neuroimaging studies. Cerebral cortex (New York, NY: 1991). 19:2767–2796.

Binney RJ, Parker GJ, Lambon Ralph MA. 2012. Convergent connectivity and graded specialization in the rostral human temporal lobe as revealed by diffusion-weighted imaging probabilistic tractography. Journal of cognitive neuroscience. 24:1998-2014.

Buckner RL, Andrews-Hanna JR, Schacter DL. 2008. The brain’s default network: anatomy, function, and relevance to disease. Ann N Y Acad Sci. 1124:1–38.

Catani M, Jones DK, ffytche DH. 2005. Perisylvian language networks of the human brain. Annals of neurology. 57:8–16.

Caverzasi E, Papinutto N, Amirbekian B, Berger MS, Henry RG. 2014. Q-ball of inferior fronto-occipital fasciculus and beyond. PloS one. 9:e100274.

Cloutman LL, Lambon Ralph MA. 2012. Connectivity-based structural and functional parcellation of the human cortex using diffusion imaging and tractography. Front Neuroanat. 6:34.

Conner CR, Ellmore TM, DiSano MA, Pieters TA, Potter AW, Tandon N. 2011. Anatomic and electro-physiologic connectivity of the language system: a combined DTI-CCEP study. Computers in biology and medicine. 41:1100–1109.

Curran EJ. 1909. A new association fiber tract in the cerebrum with remarks on the fiber tract dissection method of studying the brain. 19:645–656.

Dejerine J, Dejerine-Klumpke A. 1895. Anatomie des centres nerveux: Méthodes générales d’étude-embryologie-histogénèse et histologie. Anatomie du cerveau: Rueff.

Dick AS, Tremblay P. 2012. Beyond the arcuate fasciculus: consensus and controversy in the connectional anatomy of language. Brain: a journal of neurology. 135:3529–3550.

Donos C, Maliia MD, Mindruta I, Popa I, Ene M, Balanescu B, Ciurea A, Barborica A. 2016. A connectomics approach combining structural and effective connectivity assessed by intracranial electrical stimulation. NeuroImage. 132:344–358.

Duffau H. 2005. Intraoperative cortico-subcortical stimulations in surgery of low-grade gliomas. Expert review of neurotherapeutics. 5:473–485.

Duffau H, Gatignol P, Mandonnet E, Peruzzi P, Tzourio-Mazoyer N, Capelle L. 2005. New insights into the anatomo-functional connectivity of the semantic system: a study using cortico-subcortical electrostimulations. Brain: a journal of neurology. 128:797–810.

Duffau H, Gatignol P, Moritz-Gasser S, Mandonnet E. 2009. Is the left uncinate fasciculus essential for language? A cerebral stimulation study. Journal of neurology. 256:382–389.

Duffau H, Thiebaut de Schotten M, Mandonnet E. 2008. White matter functional connectivity as an additional landmark for dominant temporal lobectomy. Journal of neurology, neurosurgery, and psychiatry. 79:492–495.

Egger K, Yang S, Reisert M, Kaller C, Mader I, Beume L, Weiller C, Urbach H. 2015. Tractography of Association Fibers Associated with Language Processing. Clin Neuroradiol. 25 Suppl 2:231–236.

Enatsu R, Gonzalez-Martinez J, Bulacio J, Kubota Y, Mosher J, Burgess RC, Najm I, Nair DR. 2015. Connections of the limbic network: a corticocortical evoked potentials study. Cortex; a journal devoted to the study of the nervous system and behavior. 62:20–33.

Entz L, Toth E, Keller CJ, Bickel S, Groppe DM, Fabo D, Kozak LR, Eross L, Ulbert I, Mehta AD. 2014. Evoked effective connectivity of the human neocortex. Human brain mapping.

Fan L, Wang J, Zhang Y, Han W, Yu C, Jiang T. 2014. Connectivity-based parcellation of the human temporal pole using diffusion tensor imaging. Cerebral cortex (New York, NY: 1991). 24:3365–3378.

Garell PC, Bakken H, Greenlee JD, Volkov I, Reale RA, Oya H, Kawasaki H, Howard MA, Brugge JF. 2013. Functional connection between posterior superior temporal gyrus and ventrolateral prefrontal cortex in human. Cerebral cortex (New York, NY: 1991). 23:2309–2321.

Gil-Robles S, Carvallo A, Jimenez Mdel M, Gomez Caicoya A, Martinez R, Ruiz-Ocana C, Duffau H. 2013. Double dissociation between visual recognition and picture naming: a study of the visual language connectivity using tractography and brain stimulation. Neurosurgery. 72:678–686.

Glasser MF, Coalson TS, Robinson EC, Hacker CD, Harwell J, Yacoub E, Ugurbil K, Andersson J, Beckmann CF, Jenkinson M, Smith SM, Van Essen DC. 2016. A multi-modal parcellation of human cerebral cortex. Nature. 536:171–178.

Glasser MF, Rilling JK. 2008. DTI tractography of the human brain’s language pathways. Cerebral cortex (New York, NY: 1991). 18:2471-2482.

Glasser MF, Van Essen DC. 2011. Mapping human cortical areas in vivo based on myelin content as revealed by T1- and T2-weighted MRI. The Journal of neuroscience: the official journal of the Society for Neuroscience. 31:11597–11616.

Gough PM, Nobre AC, Devlin JT. 2005. Dissociating linguistic processes in the left inferior frontal cortex with transcranial magnetic stimulation. The Journal of neuroscience: the official journal of the Society for Neuroscience. 25:8010–8016.

Greenlee JD, Oya H, Kawasaki H, Volkov IO, Kaufman OP, Kovach C, Howard MA, Brugge JF. 2004. A functional connection between inferior frontal gyrus and orofacial motor cortex in human. Journal of neurophysiology. 92:1153–1164.

Greenlee JD, Oya H, Kawasaki H, Volkov IO, Severson MA, 3rd, Howard MA, 3rd, Brugge JF. 2007. Functional connections within the human inferior frontal gyrus. J Comp Neurol. 503:550–559.

Hagoort P. 2005. On Broca, brain, and binding: a new framework. Trends in cognitive sciences. 9:416–423.

Hau J, Sarubbo S, Houde JC, Corsini F, Girard G, Deledalle C, Crivello F, Zago L, Mellet E, Jobard G, Joliot M, Mazoyer B, Tzourio-Mazoyer N, Descoteaux M, Petit L. 2016. Revisiting the human uncinate fasciculus, its subcomponents and asymmetries with stem-based tractography and microdissection validation. Brain structure & function.

Hau J, Sarubbo S, Perchey G, Crivello F, Zago L, Mellet E, Jobard G, Joliot M, Mazoyer BM, Tzourio-Mazoyer N, Petit L. 2016. Cortical Terminations of the Inferior Fronto-Occipital and Uncinate Fasciculi: Anatomical Stem-Based Virtual Dissection. Front Neuroanat. 10:58.

Hickok G. 2012. The cortical organization of speech processing: feedback control and predictive coding the context of a dual-stream model. J Commun Disord. 45:393–402.

Hickok G, Poeppel D. 2004. Dorsal and ventral streams: a framework for understanding aspects of the functional anatomy of language. Cognition. 92:67–99.

Hickok G, Poeppel D. 2007. The cortical organization of speech processing. Nature reviews Neuroscience. 8:393–402.

Hoffman P, Jefferies E, Lambon Ralph MA. 2010. Ventrolateral prefrontal cortex plays an executive regulation role in comprehension of abstract words: convergent neuropsychological and repetitive TMS evidence. The Journal of neuroscience: the official journal of the Society for Neuroscience. 30:15450–15456.

Humphreys GF, Hoffman P, Visser M, Binney RJ, Lambon Ralph MA. 2015. Establishing task- and modality-dependent dissociations between the semantic and default mode networks. Proceedings of the National Academy of Sciences of the United States of America. 112:7857–7862.

Hursh JB. 1939. Conduction velocity and diameter of nerve fibers. Am J Physiol. 127:131–139.

Jackson RL, Bajada CJ, Rice GE, Cloutman LL, Lambon Ralph MA. 2018. An emergent functional parcellation of the temporal cortex. NeuroImage. 170:385–399.

Jefferies E. 2013. The neural basis of semantic cognition: converging evidence from neuropsychology, neuroimaging and TMS. Cortex; a journal devoted to the study of the nervous system and behavior. 49:611–625.

Jung J, Cloutman LL, Binney RJ, Lambon Ralph MA. 2017. The structural connectivity of higher order association cortices reflects human functional brain networks. Cortex; a journal devoted to the study of the nervous system and behavior. 97:221-239.

Keller CJ, Bickel S, Entz L, Ulbert I, Milham MP, Kelly C, Mehta AD. 2011. Intrinsic functional architecture predicts electrically evoked responses in the human brain. Proceedings of the National Academy of Sciences of the United States of America. 108:10308–10313.

Keller CJ, Honey CJ, Entz L, Bickel S, Groppe DM, Toth E, Ulbert I, Lado FA, Mehta AD. 2014. Corticocortical evoked potentials reveal projectors and integrators in human brain networks. The Journal of neuroscience: the official journal of the Society for Neuroscience. 34:9152–9163.

Keller CJ, Honey CJ, Megevand P, Entz L, Ulbert I, Mehta AD. 2014. Mapping human brain networks with cortico-cortical evoked potentials. Philosophical transactions of the Royal Society of London Series B, Biological sciences. 369.

Koubeissi MZ, Lesser RP, Sinai A, Gaillard WD, Franaszczuk PJ, Crone NE. 2012. Connectivity between perisylvian and bilateral basal temporal cortices. Cerebral cortex (New York, NY: 1991). 22:918–925.

Krieger-Redwood K, Teige C, Davey J, Hymers M, Jefferies E. 2015. Conceptual control across modalities: graded specialisation for pictures and words in inferior frontal and posterior temporal cortex. Neuropsychologia. 76:92–107.

Kubota Y, Enatsu R, Gonzalez-Martinez J, Bulacio J, Mosher J, Burgess RC, Nair DR. 2013. In vivo human hippocampal cingulate connectivity: a corticocortical evoked potentials (CCEPs) study. Clinical neurophysiology: official journal of the International Federation of. 124:1547–1556.

Lacruz ME, Garcia Seoane JJ, Valentin A, Selway R, Alarcon G. 2007. Frontal and temporal functional connections of the living human brain. The European journal of neuroscience. 26:1357–1370.

Lambon Ralph MA, Jefferies E, Patterson K, Rogers TT. 2017. The neural and computational bases of semantic cognition. Nature reviews Neuroscience. 18:42–55.

Lambon Ralph MA, Sage K, Jones RW, Mayberry EJ. 2010. Coherent concepts are computed in the anterior temporal lobes. Proceedings of the National Academy of Sciences of the United States of America. 107:2717–2722.

Lawes INC, Barrick TR, Murugam V, Spierings N, Evans DR, Song M, Clark CA. 2008. Atlas-based segmentation of white matter tracts of the human brain using diffusion tensor tractography and comparison with classical dissection. NeuroImage. 39:62–79.

Liewald D, Miller R, Logothetis N, Wagner HJ, Schuz A. 2014. Distribution of axon diameters in cortical white matter: an electron-microscopic study on three human brains and a macaque. Biological cybernetics. 108:541–557.

Mars RB, Jbabdi S, Sallet J, O’Reilly JX, Croxson PL, Olivier E, Noonan MP, Bergmann C, Mitchell AS, Baxter MG, Behrens TE, Johansen-Berg H, Tomassini V, Miller KL, Rushworth MF. 2011. Diffusion-weighted imaging tractography-based parcellation of the human parietal cortex and comparison with human and macaque resting-state functional connectivity. The Journal of neuroscience: the official journal of the Society for Neuroscience. 31:4087–4100.

Martino J, Brogna C, Robles SG, Vergani F, Duffau H. 2010. Anatomic dissection of the inferior fronto-occipital fasciculus revisited in the lights of brain stimulation data. Cortex; a journal devoted to the study of the nervous system and behavior. 46:691–699.

Martino J, Vergani F, Robles SG, Duffau H. 2010. New insights into the anatomic dissection of the temporal stem with special emphasis on the inferior fronto-occipital fasciculus: implications in surgical approach to left mesiotemporal and temporoinsular structures. Neurosurgery. 66:4–12.

Matsumoto R, Kunieda T. 2018. Cortico-Cortical Evoked Potential Mapping. In: Lhatoo SD, Kahane P, Lüders HO, editors. Invasive Studies of the Human Epileptic Brain: Principles and Practice Oxford University Press p 431–452.

Matsumoto R, Kunieda T, Nair D. 2017. Single pulse electrical stimulation to probe functional and pathological connectivity in epilepsy. Seizure: the journal of the British Epilepsy Association. 44:27–36.

Matsumoto R, Nair DR, Ikeda A, Fumuro T, Lapresto E, Mikuni N, Bingaman W, Miyamoto S, Fukuyama H, Takahashi R, Najm I, Shibasaki H, Luders HO. 2012. Parieto-frontal network in humans studied by cortico-cortical evoked potential. Human brain mapping. 33:2856–2872.

Matsumoto R, Nair DR, LaPresto E, Bingaman W, Shibasaki H, Luders HO. 2007. Functional connectivity in human cortical motor system: a cortico-cortical evoked potential study. Brain: a journal of neurology. 130:181–197.

Matsumoto R, Nair DR, LaPresto E, Najm I, Bingaman W, Shibasaki H, Luders HO. 2004. Functional connectivity in the human language system: a cortico-cortical evoked potential study. Brain: a journal of neurology. 127:2316–2330.

Matsuzaki N, Juhasz C, Asano E. 2013. Cortico-cortical evoked potentials and stimulation-elicited gamma activity preferentially propagate from lower- to higher-order visual areas. Clinical neurophysiology: official journal of the International Federation of. 124:1290–1296.

Mukherjee P, Chung SW, Berman JI, Hess CP, Henry RG. 2008. Diffusion tensor MR imaging and fiber tractography: technical considerations. AJNR American journal of neuroradiology. 29:843–852.

Noonan KA, Jefferies E, Visser M, Lambon Ralph MA. 2013. Going beyond inferior prefrontal involvement in semantic control: evidence for the additional contribution of dorsal angular gyrus and posterior middle temporal cortex. Journal of cognitive neuroscience. 25:1824–1850.

O’Muircheartaigh J, Jbabdi S. 2018. Concurrent white matter bundles and grey matter networks using independent component analysis. NeuroImage. 170:296–306.

Panesar SS, Yeh FC, Jacquesson T, Hula W, Fernandez-Miranda JC. 2018. A Quantitative Tractography Study Into the Connectivity, Segmentation and Laterality of the Human Inferior Longitudinal Fasciculus. Front Neuroanat. 12:47.

Parlatini V, Radua J, Dell’Acqua F, Leslie A, Simmons A, Murphy DG, Catani M, Thiebaut de Schotten M. 2017. Functional segregation and integration within fronto-parietal networks. NeuroImage. 146:367–375.

Peltier J, Verclytte S, Delmaire C, Pruvo JP, Godefroy O, Le Gars D. 2010. Microsurgical anatomy of the temporal stem: clinical relevance and correlations with diffusion tensor imaging fiber tracking. Journal of neurosurgery. 112:1033–038.

Perrone-Bertolotti M, Kauffmann L, Pichat C, Vidal JR, Baciu M. 2017. Effective Connectivity between Ventral Occipito-Temporal and Ventral Inferior Frontal Cortex during Lexico-Semantic Processing. A Dynamic Causal Modeling Study. Frontiers in human neuroscience. 11:325.

Ribas EC, Yagmurlu K, Wen HT, Rhoton AL, Jr. 2015. Microsurgical anatomy of the inferior limiting insular sulcus and the temporal stem. Journal of neurosurgery. 122:1263–1273.

Saito T, Tamura M, Muragaki Y, Maruyama T, Kubota Y, Fukuchi S, Nitta M, Chernov M, Okamoto S, Sugiyama K, Kurisu K, Sakai KL, Okada Y, Iseki H. 2014. Intraoperative cortico-cortical evoked potentials for the evaluation of language function during brain tumor resection: initial experience with 13 cases. Journal of neurosurgery.1–12.

Sarubbo S, De Benedictis A, Maldonado IL, Basso G, Duffau H. 2013. Frontal terminations for the inferior fronto-occipital fascicle: anatomical dissection, DTI study and functional considerations on a multi-component bundle. Brain structure & function. 218:21–37.

Saur D, Kreher BW, Schnell S, Kummerer D, Kellmeyer P, Vry MS, Umarova R, Musso M, Glauche V, Abel S, Huber W, Rijntjes M, Hennig J, Weiller C. 2008. Ventral and dorsal pathways for language. Proceedings of the National Academy of Sciences of the United States of America. 105:18035–18040.

Seghier ML. 2013. The angular gyrus: multiple functions and multiple subdivisions. The Neuroscientist: a review journal bringing neurobiology, neurology and psychiatry. 19:43–61.

Shimada S, Kunii N, Kawai K, Matsuo T, Ishishita Y, Ibayashi K, Saito N. 2017. Impact of volume-conducted potential in interpretation of cortico-cortical evoked potential: Detailed analysis of high-resolution electrocorticography using two mathematical approaches. Clinical neurophysiology: official journal of the International Federation of. 128:549–557.

Spitsyna G, Warren JE, Scott SK, Turkheimer FE, Wise RJ. 2006. Converging language streams in the human temporal lobe. The Journal of neuroscience: the official journal of the Society for Neuroscience. 26:7328–7336.

Swann NC, Cai W, Conner CR, Pieters TA, Claffey MP, George JS, Aron AR, Tandon N. 2012. Roles for the pre-supplementary motor area and the right inferior frontal gyrus in stopping action: Electrophysiological responses and functional and structural connectivity. NeuroImage. 59:2860–2870.

Takayama M, Miyamoto S, Ikeda A, Mikuni N, Takahashi JB, Usui K, Satow T, Yamamoto J, Matsuhashi M, Matsumoto R, Nagamine T, Shibasaki H, Hashimoto N. 2004. Intracarotid propofol test for speech and memory dominance in man. Neurology. 63:510–515.

Thiebaut de Schotten M, Urbanski M, Batrancourt B, Levy R, Dubois B, Cerliani L, Volle E. 2016. Rostro-caudal Architecture of the Frontal Lobes in Humans. Cerebral cortex (New York, NY: 1991).

Trebaul L, Deman P, Tuyisenge V, Jedynak M, Hugues E, Rudrauf D, Bhattacharjee M, Tadel F, Chanteloup-Foret B, Saubat C, Reyes Mejia GC, Adam C, Nica A, Pail M, Dubeau F, Rheims S, Trebuchon A, Wang H, Liu S, Blauwblomme T, Garces M, De Palma L, Valentin A, Metsahonkala EL, Bouilleret V, Landre E, Szurhaj W, Hirsch E, Valton L, Rocamora R, Schulze-Bonhage A, Mindruta I, Francione S, Maillard L, Taussig D, Kahane P, David O. 2018. Probabilistic functional tractography of the human cortex revisited. NeuroImage.

Tuch DS, Reese TG, Wiegell MR, Wedeen VJ. 2003. Diffusion MRI of complex neural architecture. Neuron. 40:885–895.

Udden J, Bahlmann J. 2012. A rostro-caudal gradient of structured sequence processing in the left inferior frontal gyrus. Philosophical transactions of the Royal Society of London Series B, Biological sciences. 367:2023–2032.

Ueno T, Saito S, Rogers TT, Lambon Ralph MA. 2011. Lichtheim 2: synthesizing aphasia and the neural basis of language in a neurocomputational model of the dual dorsal-ventral language pathways. Neuron. 72:385–396.

Usami K, Milsap GW, Korzeniewska A, Collard MJ, Wang Y, Lesser RP, Anderson WS, Crone NE. 2018. Cortical Responses to Input From Distant Areas are Modulated by Local Spontaneous Alpha/Beta Oscillations. Cerebral cortex (New York, NY: 1991).

Vassal F, Schneider F, Boutet C, Jean B, Sontheimer A, Lemaire JJ. 2016. Combined DTI Tractography and Functional MRI Study of the Language Connectome in Healthy Volunteers Extensive Mapping of White Matter Fascicles and Cortical Activations. PloS one. 11:e0152614.

Wagner AD, Pare-Blagoev EJ, Clark J, Poldrack RA. 2001. Recovering meaning: left prefrontal cortex guides controlled semantic retrieval. Neuron. 31:329–338.

Walker GM, Schwartz MF, Kimberg DY, Faseyitan O, Brecher A, Dell GS, Coslett HB. 2011. Support for anterior temporal involvement in semantic error production in aphasia: new evidence from VLSM. Brain and language. 117:110–122.

Xiang HD, Fonteijn HM, Norris DG, Hagoort P. 2010. Topographical functional connectivity pattern in the perisylvian language networks. Cerebral cortex (New York, NY: 1991). 20:549–560.

Yamao Y, Matsumoto R, Kunieda T, Arakawa Y, Kobayashi K, Usami K, Shibata S, Kikuchi T, Sawamoto N, Mikuni N, Ikeda A, Fukuyama H, Miyamoto S. 2014. Intraoperative dorsal language network mapping by using single-pulse electrical stimulation. Human brain mapping.

Yamao Y, Suzuki K, Kunieda T, Matsumoto R, Arakawa Y, Nakae T, Nishida S, Inano R, Shibata S, Shimotake A, Kikuchi T, Sawamoto N, Mikuni N, Ikeda A, Fukuyama H, Miyamoto S. 2017. Clinical impact of intraoperative CCEP monitoring in evaluating the dorsal language white matter pathway. Human brain mapping. 38:1977–1991.

Zemmoura I, Blanchard E, Raynal PI, Rousselot-Denis C, Destrieux C, Velut S. 2016. How Klingler’s dissection permits exploration of brain structural connectivity? An electron microscopy study of human white matter. Brain structure & function. 221:2477–2486.

