## supplementary data for "Connectivity Gradient in the Human Left Inferior Frontal Gyrus: Intraoperative Cortico-Cortical Evoked Potential Study"

**Figure S1.** Scatter plot of CCEP latency. The onset latency (on X axis) and the peak latency (on Y axis) are plotted about all remote CCEP responses in this study (N = 1626). Note that N1 and N2 responses are both included separately even if they are found in the same waveform. The color of dot indicates the classification suggested by the cluster analysis (Ward's method). The red dots are regarded as typical N1 responses. The blue dots are typical as N2. The green dots seems to constitute another entity between N1 and N2. The bar graphs on the top and right of the scatter plot show the histogram of onset and peak. The red bars in the histogram corresponds to the cluster of typical N1 response.

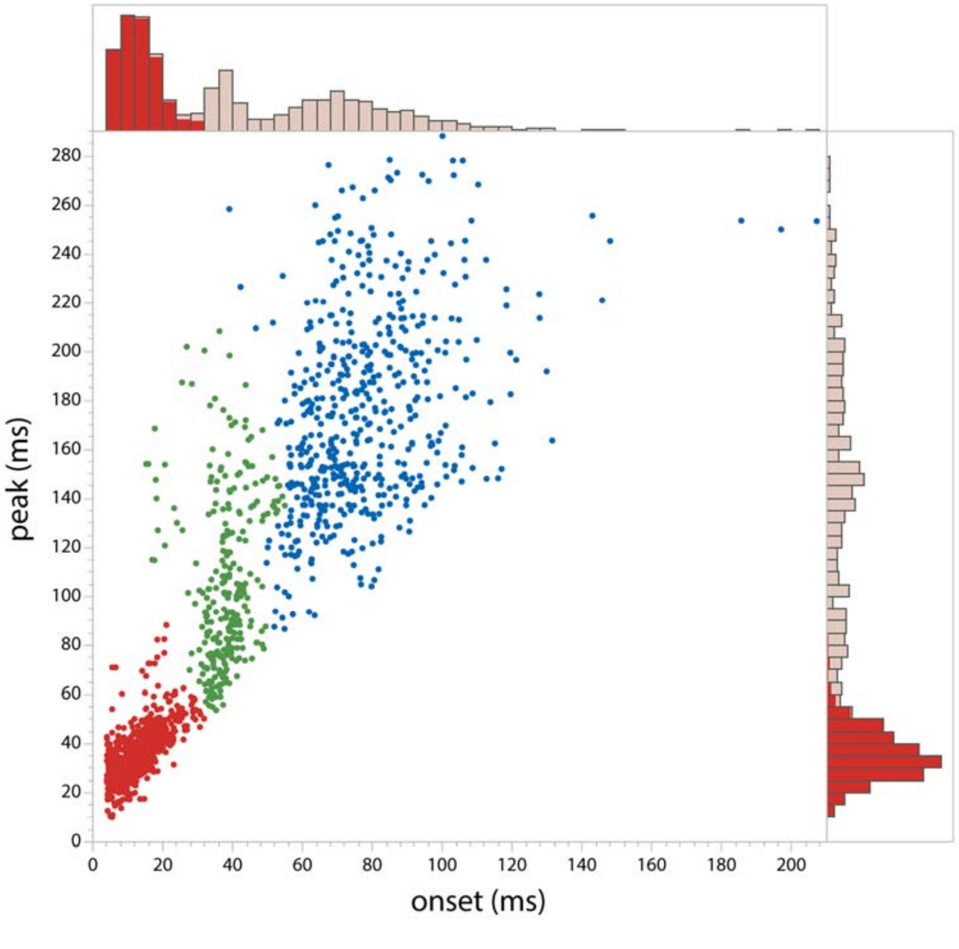

**Figure S2.** Scatter plot of the stimulus-response distance and the onset latency of the N1 response was made about all temporal response by the stimulation of IFG (N = 93). Red line indicates the regression line for the stimulation of all parts in IFG ($p < 0.05$). No regression line could be drawn for each part in IFG.

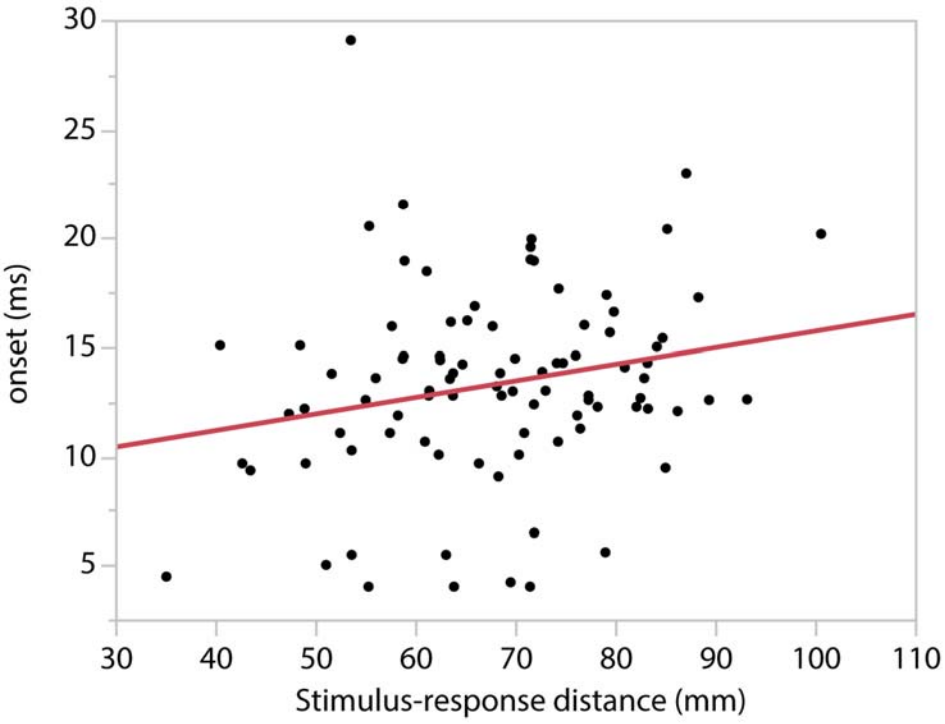

**Movie. “4D CCEP map”.**

Each movie represents the time-sequence of CCEP response by the stimulation of each part in IFG (pOrb, pTri or pOpe). To make a movie, we created images by 1 ms from 30 ms before the stimulus to 300 ms after the stimulus (total 331 images). Each image was derived from the standard brain surface painted according to the CCEP response distribution. The technical details in making the CCEP response distribution are described in another supplementary material. To visualize the volume data on the brain surface model, we painted each vertex by the intensity of the nearest voxel according to the color scale, skipping those voxels which are covered by less than 5 subjects within 15 mm. Colored spheres indicates the location of the stimulus sites (red: IFG pOrb, orange: IFG pTri, green: IFG pOpe).

**Technical details in making the CCEP response distribution**

We created the time-sequence of 3D volume image that represents the time course of the electric potential averaged across all subjects, which is called "averaged response map" hereafter. Averaging across subject is not simple because the location of electrodes are different. To solve that, we first accumulated the electric potential at all electrodes in the MNI space (voxel size: 2mm, isometric) through all subjects, plotting each recorded value (electric potential) at the voxel of electrode location. Then, we smoothed the volume (accumulated potential data) with Gaussian kernel (FWHM 6 mm, kernel size 10 mm) to absorb the inherent dispersion in the electrode determination. We set FWHM as twice as much as the electrode diameter (3 mm), for we thought it is the lowest (humblest) precision when electrodes location are determined by the intraoperative picture. In this way, we obtain the volume image of the accumulation of recorded potentials which we call the "accumulated potential map". Similarly, we obtain the "observation density map" by plotting "1" instead of the electric potential in each electrode. Note that we use all recorded data if the subject has several stimulus sites in the concerned area (pOrb, pTri or pOpe). Finally, we divide the "accumulated potential map" by the "observation density map" to obtain the "averaged response map". The calculation is skipped where the observation density is zero. We applied the spatial smoothing with Gaussian filter

(FWHM 10 mm) to the “averaged response map”. The electrode interval, 10 mm, seems suitable for FWHM to make the output image close to what we see in the rough estimation of the spatial distribution.

**Technical details of reciprocality analysis**

A judgement of reciprocality was made from the CCEP database in the following three steps, which were implemented by the in-house MATLAB script. We checked the reciprocality of fronto-temporal and fronto-parietal connections separately. Taking the fronto-temporal connections as an example, our detailed method was as follows:

1. We extracted the temporal responses by frontal stimulation (= IFG pOrb stimulation in this study) from the CCEP database. We regard them as "anterograde" connections because we are going to search retrograde connectivity and to judge their reciprocality. The electrodes with response were searched in the CCEP database in two ways; one is all of the responses and the other is the max responses. For both type of response, we calculated the reciprocality rate. For each fronto-temporal connection, we repeated the below (step 2 and 3) to judge whether it is reciprocal or not.
2. We searched for frontal responses by the stimulation through the temporal response electrodes, which we call "retrograde" stimulation, For all the temporal response electrodes identified in the previous step, we checked whether a stimulus through the temporal response electrode was performed or not and whether it caused a CCEP response in the frontal stimulus electrode or not. Note that we could not stimulate through all of the temporal response electrodes due to the limited time in the

operating room. Instead, we focused on the temporal response electrodes actually examined and calculated the rate of the reciprocality among them.

3. We made a judgement of reciprocality. When the retrograde stimulation evoked a max response in at least one of the paired stimulus electrodes in the frontal lobe, we considered the frontal stimulus site has a reciprocal connection with the temporal response site.

Through the three steps, we assessed the rate of reciprocal connection. We assessed reciprocality in six groups stratified by area (fronto-temporal vs. fronto-parietal) and the type of anterograde response (max response, any response, or no response). We calculated the reciprocality rate not only for the response electrodes but for the no-response electrodes (i.e., those which did not show any response by frontal stimulation) to obtain the negative controls.
